# G9a and Sirtuin6 epigenetically modulate host cholesterol accumulation to facilitate mycobacterial survival

**DOI:** 10.1101/2021.02.27.433201

**Authors:** Praveen Prakhar, Bharat Bhatt, Tanushree Mukherjee, Gaurav Kumar Lohia, Ullas Kolthur-Seetharam, Nagalingam Ravi Sundaresan, R.S. Rajmani, Kithiganahalli Narayanaswamy Balaji

**Affiliations:** Department of Microbiology and Cell Biology, Indian Institute of Science, Bangalore – 560012, Karnataka, India; Department of Biological Sciences, Tata Institute of Fundamental Research, Mumbai – 400005, Maharashtra, India; Centre for Infectious Disease Research, Indian Institute of Science, Bangalore – 560012, Karnataka, India

**Keywords:** *Mycobacterium tuberculosis*, epigenetic modifications, G9a, SIRT6, cholesterol accumulation

## Abstract

Cholesterol derived from the host milieu forms a critical factor for mycobacterial pathogenesis. However, the molecular circuitry co-opted by *Mycobacterium tuberculosis* (Mtb) to accumulate cholesterol in host cells remains obscure. Here, we report that a functional amalgamation of WNT-responsive histone modifiers G9a (H3K9 methyltransferase) and Sirt6 (H3K9 deacetylase) orchestrate cholesterol build-up in *in-vitro* and *in-vivo* models of Mtb infection. Mechanistically, G9a, along with SREBP2, drives the expression of cholesterol biosynthesis and uptake genes; while Sirt6 represses the genes involved in cholesterol efflux. The accumulated cholesterol promotes the expression of antioxidant genes leading to reduced oxidative stress, thereby supporting Mtb survival. In corroboration, loss-of-function of G9a *in vitro* and *in vivo* by pharmacological inhibition; or utilization of BMDMs derived from *Sirt6* KO mice or *in vivo* infection in *Sirt6* heterozygous mice; hampers host cholesterol accumulation and restricts Mtb burden. These findings shed light on the novel roles of G9a and Sirt6 during Mtb infection and highlight the previously unknown contribution of host cholesterol in potentiating anti-oxidative responses for aiding Mtb survival.

## Introduction

*Mycobacterium tuberculosis* (Mtb) rewires host cellular machinery to subvert protective immune responses and achieve a secure and nutrient-rich niche. Emerging evidences highlight the implication of epigenetic factors in Mtb-driven tuning of gene expression to effectuate such immune evasion^1–3^. Reports suggest that the histone methyltransferase (HMT) EZH2 epigenetically down-modulates MHC-II presentation^4^; SET8 HMT governs immune processes such as apoptosis, oxidative stress and cytokine secretion^5^; while certain mycobacterial proteins themselves gain access to host chromatin and modulate a plenitude of immune genes^6^. One of the classical features of tuberculosis (TB) infection involves the accumulation of neutral lipids, cholesterol and cholesteryl esters to generate foamy macrophage (FM) phenotype^7,8^. Although, certain studies report that lipid droplets do not serve as an important source of nutrient for Mtb, hence, do not affect Mtb growth^9^ and inhibition of fatty acid oxidation restricts intracellular growth of Mtb via ROS production^10^; there are evidences to support that the formation of FMs positively correlates with mycobacterial virulence and the loss of lipids from these cells compromises mycobacterial survival by not only limiting nutrients but also by curbing the requisite cues for altering hosts’ ER stress, survival pathways and autophagy levels^11–19^. In this context, cholesterol serves essential functions for Mtb in the acquisition of dormancy and resistance to antibiotics in both, *in vitro* and *in vivo* systems^14,20^. To our interest, it has been recently reported that host cells such as macrophages form a major source of cholesterol for intracellular Mtb ^21^ and also possibly for extracellular Mtb released into the caseated or cavitated TB granuloma lesions^14^. However, information regarding the mechanisms regulating cholesterol accumulation in hosts during Mtb infection requires extensive investigation. Existing literature suggests that genes of cholesterol biosynthesis and homeostasis are epigenetically governed by miRNAs (miR-33a, miR-185), histone deacetylases (HDAC3, Sirt2, Sirt6) and HMTs (G9a) under distinct conditions^22,23^. Amongst these, the study on Sirt2 does not correlate its *in vivo* activity with chronic Mtb infection^24^. Contrastingly, activation of the nuclear Sirtuin, Sirt1, has recently been ascribed with restriction of Mtb growth by augmenting autophagy^25^. Sirt1 has also been shown to be involved in lipid metabolism, stress response, anti-inflammatory response and cellular senescence in diverse contexts^26–30^. However, the contribution of the other nuclear Sirtuin, i.e. Sirt6, during infections, specifically mycobacterial infection, has not been addressed so far; although it has been enlisted as an upregulated gene in Mtb infection-related transcriptomic dataset^31^. Sirt6 majorly confers H3K9- and H3K56-deacetylation and is known to be associated with life span, genome stability and tumorigenesis^32–35^. Strikingly, Sirt6 has been identified as a potential regulator of SREBP1/2 functions, a major transcription factor for cholesterol metabolism^36^. We were piqued to unravel the epigenetic contribution of Sirt6 in accumulating cholesterol during TB infection. Additionally, evidences from the literature suggest that apart from acetylation, methylation of H3K9 (mono- and di-), conferred by G9a, imparts crucial epigenetic signatures for shaping immunological fates during various pathophysiological conditions, such as T cell differentiation, immunological memory, viral latency and endotoxin tolerance^37–42^. With this premise, we focused on elucidating the interplay of H3K9 methylation and acetylation by G9a and Sirt6, respectively, in defining cholesterol accumulation during TB infection.

We find that Mtb induces the production of G9a and Sirt6, which contribute to epigenetically driven differential expression of cholesterol biosynthesis, uptake and efflux genes, thereby allowing cholesterol accumulation during infection. Mechanistically, WNT signaling was found to govern the levels of G9a and Sirt6 upon Mtb infection. WNT signaling has earlier been implicated in cell proliferation, migration, immunological processes and also in shaping immune responses during Further, the accumulated cholesterol was found to aid mycobacterial survival by promoting anti-oxidative factors. Loss-of-function of G9a using a pharmacological inhibitor and that of Sirt6 using *Sirt6* heterozygous mice in an *in vivo* mouse TB model was found to hamper host cholesterol accumulation and restrict Mtb burden. This was also corroborated by lung histopathology, which indicated a reduced severity of TB in mice lacking G9a and Sirt6 functions. Together, we show for the first time that G9a and Sirt6 are upregulated during Mtb infection; and in conjunction mediate TB pathogenesis by epigenetically reprogramming cholesterol accumulation. Besides, this study underscores the relevance of specific G9a and Sirt6 inhibitors as plausible anti-TB adjuvants.

## Results

### Interception of G9a and Sirt6 leads to restricted mycobacterial burden

Infection of host macrophages with pathogenic Mtb H37Rv was found to induce the expression of the HMT G9a (encoded by *Ehmt2*) at transcript as well as protein level; without any significant change in the global H3K9 monomethylation pattern (**Fig. 1A; S1A, B**). Alongside, we report that Mtb H37Rv augments the expression of the HDAC Sirt6. However, the corresponding global H3K9 acetylation mark also remains unaltered (**Fig. 1A; S1A, B**). Importantly, we corroborated our findings in human primary macrophages (**Fig. 1B; S1C**) as well as in a mouse model of pulmonary TB infection (**Fig. 1C, D; S1D**), wherein, we observed an enhanced expression of the concerned epigenetic factors at the transcript and protein level during Mtb H37Rv infection. Further, the induction of G9a and Sirt6 was found to be specific to virulent species of mycobacterium as infection of mouse peritoneal macrophages with the non-pathogenic *Mycobacterium smegmatis* showed only a weak expression of G9a and Sirt6 (**Fig. S1E**). To evaluate the possible contribution of these epigenetic factors to Mtb infection, *in vitro* CFU assays were performed. Inhibition of G9a using a specific pharmacological inhibitor (BIX-01294) was found to compromise Mtb H37Rv burden after 48 h of *in vitro* infection (**Fig. 1E**). Next, mycobacterial CFU was assessed in BMDMs derived from WT (littermate control) and *Sirt6* knockout (KO) mice as the premature ageing and death of Sirt6 KO mice by 4 weeks post-natally^45^ hinders the isolation of thioglycolate-elicited peritoneal macrophages and long-term *in vivo* experiments. Sirt6 expression was assessed in the lung homogenate of WT, Sirt6 heterozygous and Sirt6 KO mice **(Fig S1F)**. We found the Mtb H37Rv burden to be restricted in *Sirt6* KO BMDMs (**Fig. 1F**). Further, *in vitro* silencing of *Ehmt2* or *Sirt6* individually lead to a compromised mycobacterial CFU, that was further diminished in mouse peritoneal macrophages knocked down for both *Ehmt2* and *Sirt6* (**Fig. 1G; Fig. S1G**: knockdown validation; **Fig. S1H**: cell viability assessment). These results suggest a critical role for the epigenetic modifiers G9a and Sirt6 in the successful pathogenesis of mycobacterium.

**Figure 1.**
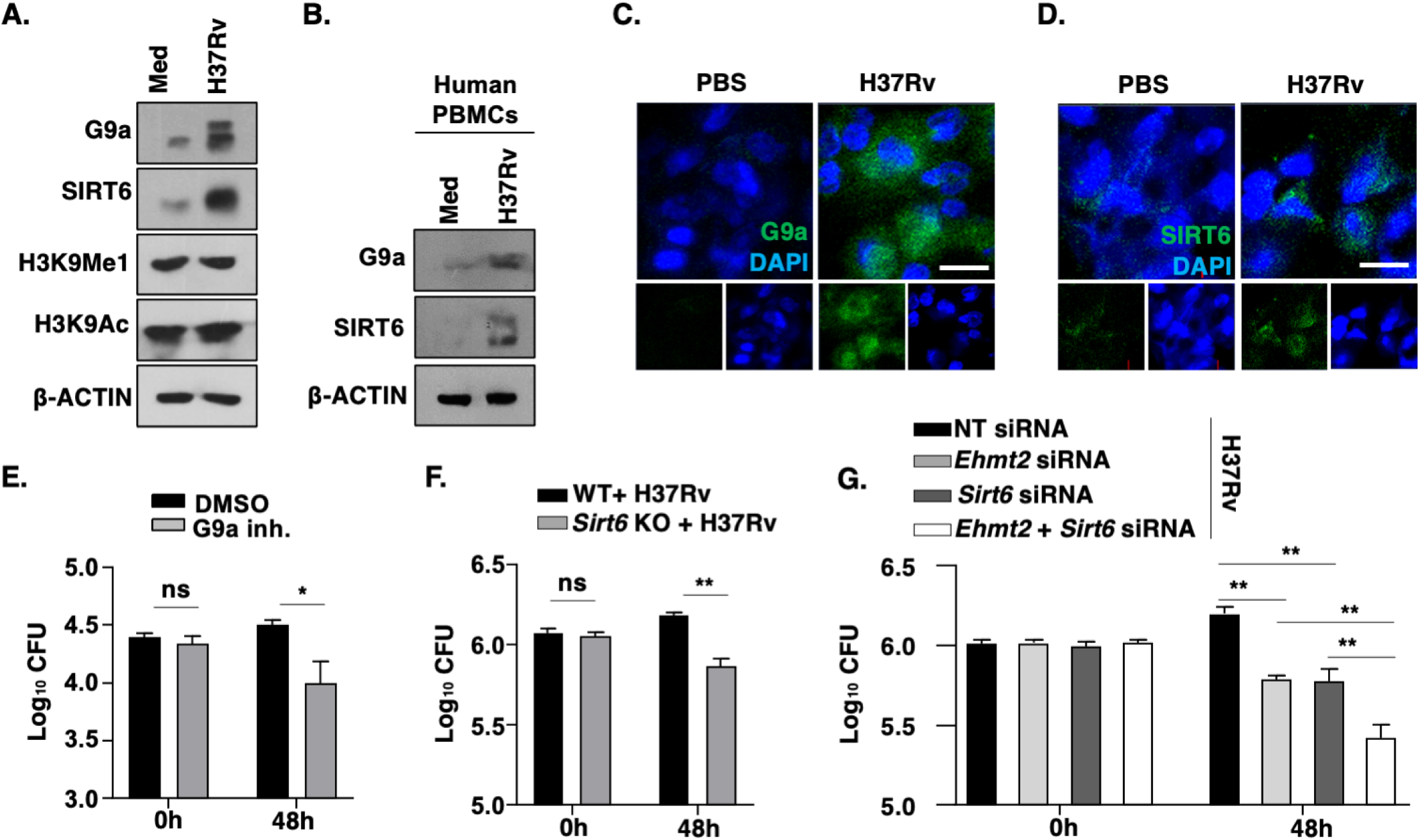
Interception of G9a and Sirt6 leads to restricted mycobacterial burden. **(A)** BALB/c peritoneal macrophages were infected with H37Rv for 12 h, protein level of G9a, SIRT6 and their respective histone modification marks, H3K9me1 and H3K9Ac, were assessed by immunoblotting. **(B**) Immunoblot for G9a and SIRT6 in human PBMCs infected with H37Rv for 12 h. **(C, D)** *In vivo* expression of G9a **(C)** and SIRT6 **(D)** was analyzed in lung cryosections of uninfected mice and mice infected with H37Rv for 56 days by immunofluorescence. **(E-G)** *In vitro* CFU was assessed 48 h post H37Rv infection under the following conditions: **(E)** in BALB/c mouse peritoneal macrophages treated with G9a inhibitor (5 μM) or **(F)** in BMDMs derived from WT (littermate control) or *Sirt6* KO mice or **(G)** in BALB/c mouse peritoneal macrophages transiently transfected with siRNAs against G9a or Sirt6 or both. The MOI of infection is 1:10 (macrophage:mycobacteria) for all the *in vitro* experiments. All data represents the mean ± SEM from 3 independent experiments; *, P < 0.05, **, P < 0.01; ns, not significant (Student’s t-test for E-F and One-way ANOVA for G) and the blots are representative of 3 independent experiments. Med, medium (uninfected/untreated cells maintained in DMEM supplemented with 10% heat inactivated FBS for the entire duration of the experiment); WT, wild type; KO, knock out; inh., inhibitor; NT, non-targeting; BMDM, bone marrow derived macrophages; PBMC, peripheral blood mononuclear cells. Scale bar, 10μm.

### G9a and Sirt6 effectuate cholesterol accumulation during mycobacterial pathogenesis

As described earlier, among various factors, host-derived cholesterol forms an integral part of mycobacterial pathogenesis *in vitro* and *in vivo*. In this context, virulent Mtb H37Rv infection was found to trigger cholesterol accumulation in host macrophages, unlike *M. smegmatis* infection, as assessed by Filipin staining (**Fig. S2A, B**). The same was mirrored in the lungs of mice infected with Mtb H37Rv, wherein staining lung cryosections with Filipin showed a significant increase in cholesterol accumulation specifically in macrophages (**Fig. S2C, D**). With the premise that both G9a and Sirt6 have been reported to epigenetically regulate cholesterol levels^22^, we sought to explore their contribution to cholesterol accumulation in the context of Mtb infection. It was observed that *in vitro* silencing of *Ehmt2* and *Sirt6* via specific siRNAs led to a marked decline in the ability of Mtb H37Rv to furnish cholesterol accretion (**Fig. 2A, B**). Further, significantly less cholesterol was detected by Filipin staining in BMDMs derived from *Sirt6* KO mice, even in the presence of Mtb H37Rv infection (**Fig. 2C, D**). Substantiating the same, macrophage-specific accumulation of cholesterol was reduced in the lungs of G9a inhibitor-treated mice (**Fig. 2E, F**) and *Sirt6* heterozygous mice (**Fig. 2G, H**). These evidences strongly indicate the ability of Mtb to utilize G9a and Sirt6 for mediating the essential process of cholesterol accumulation in host cells.

**Figure 2.**
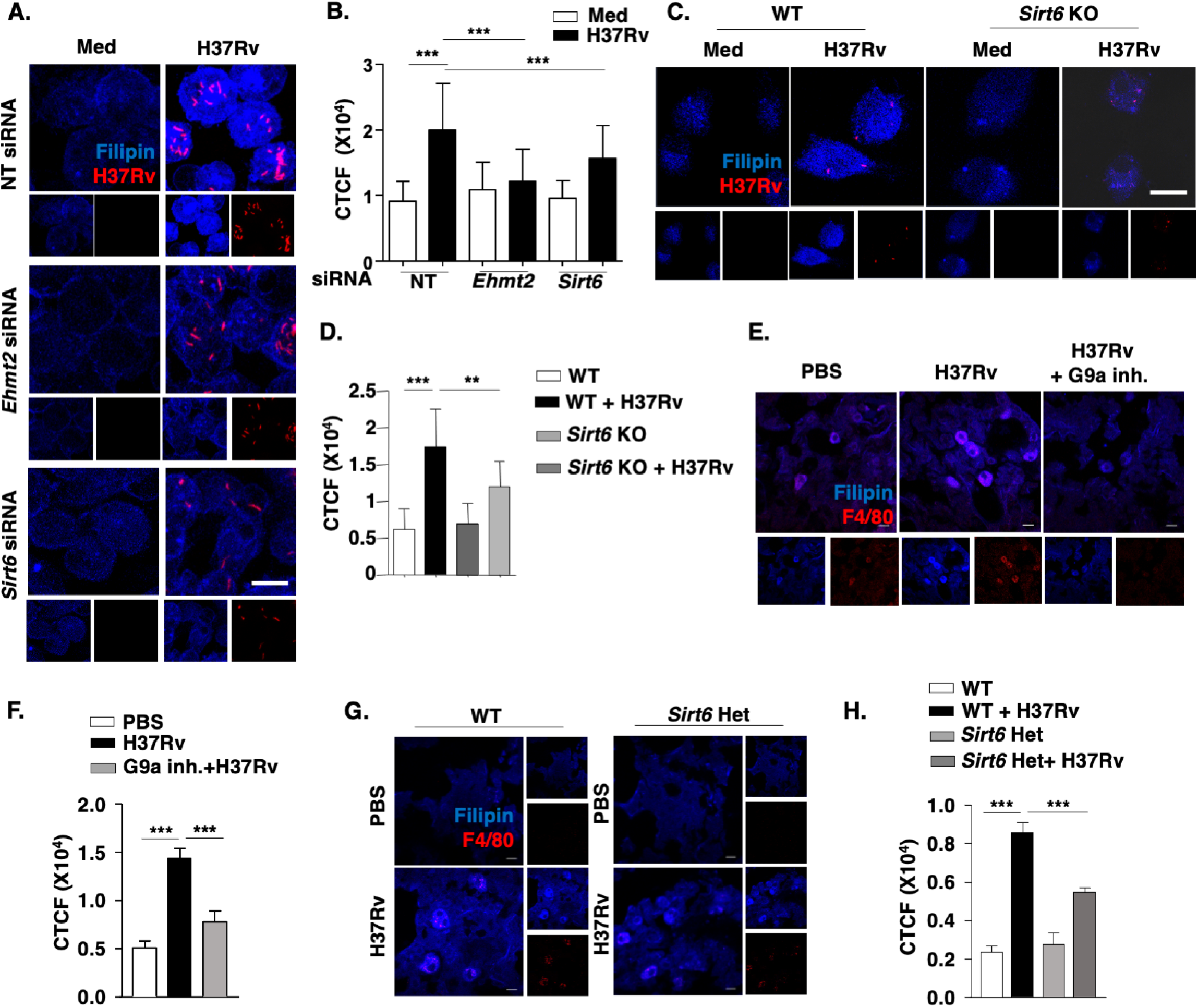
G9a and Sirt6 induce cholesterol accumulation during mycobacterial pathogenesis. **(A, B)** BALB/c mouse peritoneal macrophages transfected with NT or *Ehmt2* or *Sirt6* siRNA were assessed for free cholesterol level upon infection with tdTomato-expressing H37Rv for 48 h by immunofluorescence. **(A)** Representative images of Filipin stained macrophages and **(B)** its quantification (n=200-300). **(C, D)** BMDMs from WT (littermate control) and *Sirt6* KO mice were utilized to assess free cholesterol by Filipin staining upon tdTomato-expressing H37Rv infection for 48 h. **(C)** Representative images and **(D)** its quantification (n=200-300). **(E, F)** Lung cryosections from uninfected or 56 days H37Rv-infected/ G9a inhibitor (40mg/kg) treated BALB/c mice were assessed for free cholesterol by Filipin staining in macrophages stained with F4/80: **(E)** representative images and **(F)** quantification of Filipin staining in F4/80 positive cells (i.e. macrophages). **(G, H)** Lung cryosections of uninfected and infected WT (littermate control) and *Sirt6* het mice were assessed for free cholesterol levels by Filipin staining in macrophages stained by F4/80: **(G)** representative images and **(H)** quantification of Filipin staining in F4/80 positive cells (i.e. macrophages). *In vivo* data represents the mean ± SEM from 2-3 mice. The MOI of infection is 1:10 (macrophage: mycobacteria) for all the *in vitro* experiments. All data represents the mean ± SEM from 3 independent experiments, *, P < 0.05; **, P < 0.01; ***, P < 0.001 (One-way ANOVA for B, D, F and H). Med, medium; WT, wild type; KO, knock out; het, heterozygous; inh., inhibitor; MFI, mean fluorescence intensity; CTCF, corrected total cell fluorescence; F4/80, macrophage marker; BMDM, bone marrow derived macrophages. Scale bar, 10μm.

### Cholesterol biosynthesis and transport genes are differentially regulated by G9a and Sirt6

The accumulation of cholesterol in a given cell or tissue results from the coordinated interplay of genes involved in its biosynthesis and uptake on one hand, and its efflux on the other. To this end, the status of the pertinent markers (23 genes) of each function was assessed for their transcript level expression during infection with Mtb H37Rv *in vitro* and *in vivo* (**Fig. S2E, F**). We observed that genes involved in cholesterol uptake (*Lrp2*) and biosynthesis (*Aacs*, *Hmgcs1*, *Mvd*, *Dhcr24*) were significantly upregulated during infection; while those implicated in efflux (*Abca1*, *Abcg1*) showed a marked downregulation. We further assessed the transcript levels of the altered genes in human PBMCs infected with Mtb H37Rv. Corroborating our *in vitro* and *in vivo* mice data, we observed a significant increase in the transcript levels of genes involved in cholesterol uptake (*Lrp2*) and biosynthesis (*Aacs*, *Hmgcs1*, *Mvd*, *Dhcr24*) with a downregulation in the expression of efflux genes (*Abca1*, *Abcg1*) in H37Rv-infected human PBMCs (**Fig. S2G**). Interestingly, this differential gene expression was found to be finely tuned by the combined activities of G9a and Sirt6. While depletion of G9a function by siRNA-mediated knock-down compromised the expression of the biosynthesis and uptake genes (*Lrp2, Aacs, Hmgcs, Mvd, Dhcr24*) at the transcript level (**Fig. 3A**); that of Sirt6 rescued the Mtb-dependent downregulation of cholesterol efflux genes (*Abca1, Abcg1*) (**Fig. 3B**). Inline, overexpressing Sirt6 led to a marked decrease in *Abca1* and *Abcg1* expression (**Fig. S3A**). The transcript level profiling performed for the concerned genes in the lungs of mice treated with G9a inhibitor *in vivo* or *Sirt6* heterozygous mice also yielded a similar pattern (**Fig. 3C, D**). These findings were validated at the protein level, where G9a inhibitor limited the surface expression of LRP2 during infection, both *in vitro* and *in vivo* (**Fig. 3E, F**); and ABCA1 protein expression was found to be elevated in the lungs of Mtb H37Rv-infected *Sirt6* heterozygous mice (**Fig. 3G**), compared to that in the infected wild type controls. The protein level of ABCA1 was also rescued in macrophages knocked down for Sirt6 (**Fig. S3B**). The current set of observations prompted us to delineate the G9a- and Sirt6-driven mechanism of differential regulation of cholesterol biosynthesis, uptake and efflux genes.

**Figure 3.**
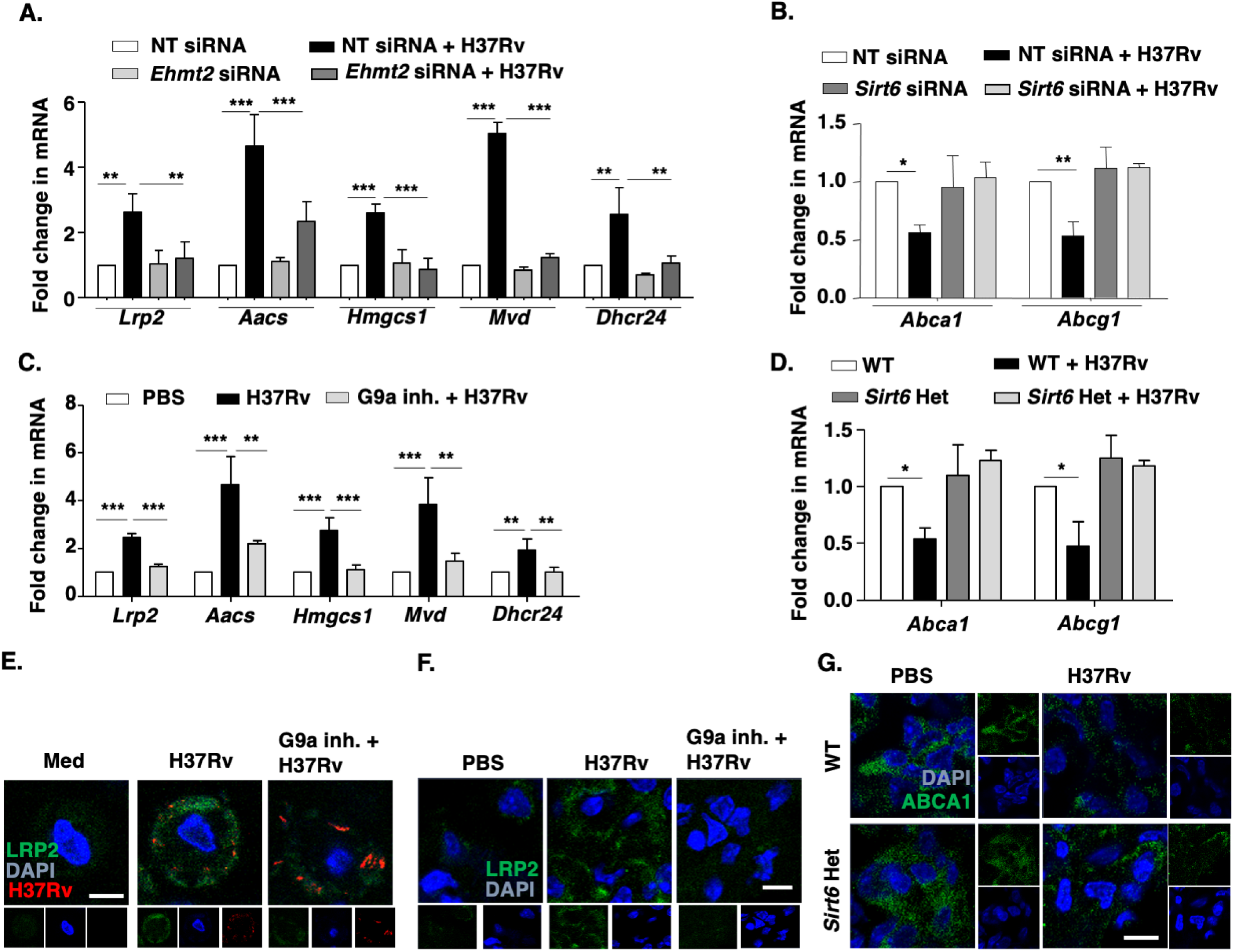
Cholesterol biosynthesis and transport genes are selectively regulated by G9a and Sirt6. **(A, B)** BALB/c mouse peritoneal macrophages were transfected with NT or *Ehmt2* or *Sirt6* siRNA. Transfected cells were infected with H37Rv for 12 h and the expression of the indicated genes was assessed by qRT-PCR. **(C, D)** RNA was isolated from the lung homogenates from the indicated groups of mice after 56 days of total infection and the transcript levels of the indicated cholesterol metabolism genes was analyzed by qRT-PCR. **(E)** Surface expression of LRP2 was analyzed by immunofluorescence in BALB/c peritoneal macrophages pre-treated with G9a specific inhibitor (5μM) for 1 h followed by infection with tdTomato-expressing H37Rv for 12 h. **(F)** Lung cryosections from uninfected or 56 days H37Rv-infected/ G9a inhibitor (40mg/kg) treated BALB/c mice were assessed for surface expression of LRP2 by immunofluorescence. **(G)** Lung cryosections of uninfected and 56 days H37Rv-infected WT (littermate control) and *Sirt6* het mice were assessed for the protein level of ABCA1 by immunofluorescence. *In vivo* data represents the mean ± SEM from 2-3 mice. The MOI of infection is 1:10 (macrophage: mycobacteria) for all the *in vitro* experiments. All data represents the mean ± SEM from 3 independent experiments, *, P < 0.05; **, P < 0.01; ***, P < 0.001 (One-way ANOVA for A-D). Med, medium; WT, wild type; het, heterozygous; inh., inhibitor; NT, non-targeting. Scale bar, 10μm.

### G9a-SREBP2 and Sirt6 fine-tune cholesterol accumulation during Mtb infection

The transcription factor SREBP2 (encoded by *Srebf2*) is a well-established master regulator of cholesterol biosynthesis genes. However, it functions in tight association with accessory transcription activators and regulators^46^. We hypothesized the possibility of SREBP2 and G9a to interact and together bring about the augmented expression of cholesterol biosynthesis and uptake genes. In this regard, first we show that Mtb H37Rv induces SREBP2 expression and immunopulldown analysis indicates that SREBP2 interacts with G9a during Mtb H37Rv infection (**Fig. 4A**). Loss-of-function of SREBP2 using specific siRNA (**Fig. S3C**, knockdown validation) compromised the expression of positive factors of cholesterol accumulation (*Lrp2, Aacs, Hmgcs, Mvd, Dhcr24*) (**Fig. 4B**). In line with this, Mtb H37Rv burden was significantly restricted in macrophages knocked down for *Srebf2 in vitro* (**Fig. 4C**). Next, we verified the role of G9a HMT at the chromatin level by assessing the recruitment of G9a to SREBP2 binding sites at the promoters of cholesterol biosynthesis and uptake genes. Both, G9a occupancy and associated H3K9me1 marks were found to be elevated during Mtb H37Rv infection (**Fig. 4D, E**). Further, sequential ChIP analysis showed an enhanced co-occupancy of the concerned promoters with G9a and SREBP2 (**Fig. 4F**), thus confirming the concerted action of SREBP2 and G9a in driving the expression of cholesterol biosynthesis and uptake genes (*Lrp2, Aacs, Hmgcs, Mvd, Dhcr24*). Contrary to this, Sirt6 was found to occupy the promoters of *Abca1* and *Abcg1* during Mtb H37Rv infection, leading to concomitantly decreased acetylation marks; and supporting the initially observed downregulation of the cholesterol efflux genes during Mtb infection (**Fig. 4G, H**). Further, since H3K9me2 conferred by G9a renders a closed chromatin state and transcriptional downregulation, the contribution of G9a in the reduced expression of ABCA1 was analyzed. It was found that mycobacteria lost the ability to downregulate ABCA1 in G9a knocked down macrophages (**Fig. S3D, E**); indicating the partial dependence of cholesterol efflux genes on the repressive function of G9a. To understand this interplay of G9a and Sirt6, a time kinetics ChIP assay was performed. Interestingly, it was observed that the temporal recruitment of Sirt6 and G9a allows the repression of cholesterol efflux genes by both Sirt6 and G9a at early time points (12 h) (**Fig. S4A, B**), that is then sustained only by G9a at later times of infection (24 h) (**Fig. S4C**). However, unlike for the biosynthesis/uptake genes, SREBP2 was not found to be involved in the regulation of the efflux genes (**Fig. S4D**). Also, the cholesterol biosynthesis and uptake genes (*Lrp2, Aacs, Hmgcs, Mvd, Dhcr24*) were only regulated by G9a and were found to be independent of Sirt6 (**Fig. S4E-G**).

**Figure 4.**
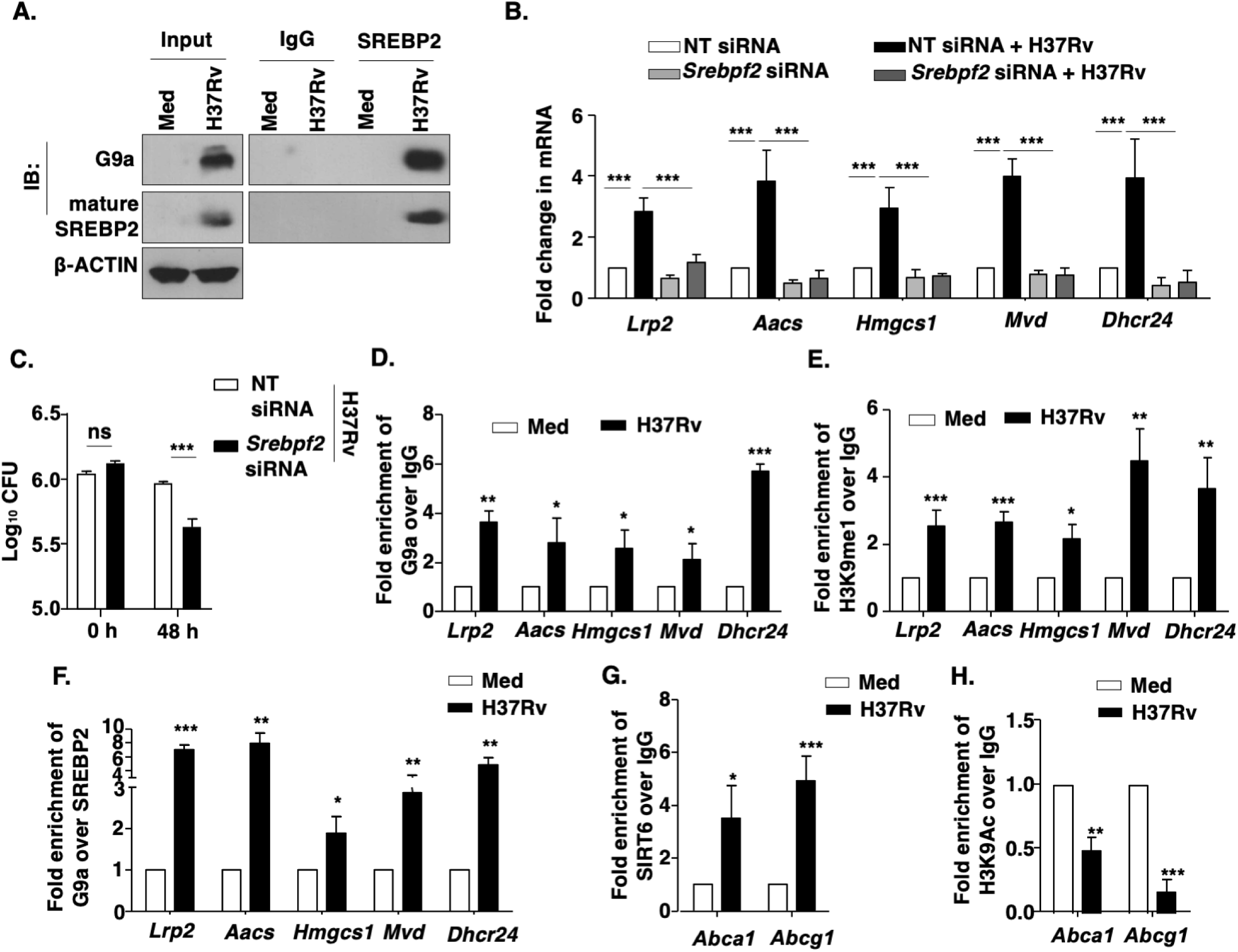
Mtb-induced G9a and Sirt6 mediate fine-tuning of cholesterol accumulation. **(A)** SREBP2 was immunoprecipitated in whole cell lysates of BALB/c mouse peritoneal macrophages infected with H37Rv for 12 h to assess its interaction with G9a by immunoblotting. **(B, C)** BALB/c mouse peritoneal macrophages were transfected with *Srebpf2* siRNA and **(B)** infected for 12 h with H37Rv to assess the expression of cholesterol accumulation genes by qRT-PCR, or **(C)** infected with H37Rv for 48 h to analyze the *in vitro* CFU. **(D, E)** ChIP assay was performed to affirm the **(D)** recruitment of G9a and **(E)** corresponding H3K9me1 mark, on the promoters of the indicated genes upon 12 h infection with H37Rv in BALB/c mouse peritoneal macrophages. **(F)** Sequential ChIP was conducted to assess the co-recruitment of SREBP2 and G9a at the promoters of *Lrp2*, *Aacs*, *Hmgcs1*, *Mvd* and *Dhcr24* in mouse peritoneal macrophages infected with H37Rv for 12 h. **(G, H)** BALB/c mouse peritoneal macrophages infected with H37Rv for 12 h were analyzed by ChIP for **(G)** SIRT6 recruitment and **(H)** H3K9Ac mark, on the promoters of *Abca1* and *Abcg1*. The MOI of infection is 1:10 (macrophage: mycobacteria) for all the *in vitro* experiments. All data represent the mean ± SEM from 3 independent experiments. The blots are representative of 3 independent experiments. *, P < 0.05; **, P < 0.01; ***, P < 0.001; ns, not significant (One-way ANOVA for B, Student’s t-test for C-H). Med, Medium; NT, non-targeting.

### Cholesterol accumulation mitigates oxidative stress during mycobacterial infection

The orchestrated accumulation of cholesterol by Mtb-induced G9a and Sirt6 provides insights into the crucial functions that cholesterol might effectuate to favor Mtb survival. As discussed, the contribution of cholesterol as a source of nutrition for mycobacteria is widely accepted. However, evidences from the literature suggest numerous alternate implications of cholesterol in cellular homeostasis and responses to stimuli^47,48^. Moreover, supplementation of exogenous cholesterol helped in mitigating the toxic effects of bile acid and lipids in Nonalcoholic steatohepatitis by enhancing the expression of NRF2 and HIF-1α^49^. Similarly, Cholesterol crystals present in the atherosclerotic plaques also act as a stimulus to regulate Nrf-2^50^. We report that mycobacteria-induced cholesterol accumulation renders the expression of a principal transcription activator of antioxidant genes, NRF2 (encoded by *Nfe2l2*) that then leads to the expression of its target genes (**Fig. S5A-D**). In this line, perturbation of G9a or Sirt6 using specific siRNAs compromised the expression of NRF2-target genes *Trxrd1*, *Nqo1*, *Hmox1*, *Gsr*, *Gpx1*, *Sod1* at the transcript level (**Fig. S5E**) and protein level (**Fig. 5A**). We next verified that the observed loss of antioxidant gene expression indeed resulted from hampered accumulation of cholesterol in G9a- and Sirt6-knocked down cells. To this end, first we utilized siRNAs against the five G9a-dependent genes found to be essential for cholesterol biosynthesis and uptake (*Lrp2, Aacs, Hmgcs, Mvd, Dhcr24*); and verified the abrogation of cholesterol accumulation during Mtb H37Rv infection to result from the specific downregulation of the five concerned targets (**Fig. S6A-C**), while negating the implication of any other Mtb-unresponsive cholesterol biosynthesis genes in the process (**Fig. S6D**). In these cholesterol deficient cells, we found a significant reduction in the expression of antioxidant genes at the transcript and protein level (**Fig. S5E, 5B**); implicating cholesterol in driving antioxidant gene expression. Consequently, increased oxidative stress was observed in G9a and Sirt6 knocked down cells (**Fig 5C-D**) as well as in cells deficient of G9a-dependent cholesterol biosynthesis and uptake gene expression (**Fig. 5E-F**). Corollary to this, we also observed that *in vitro* depletion of cholesterol accumulation genes (**Fig. 5G**) or *Nfe2l2* (**Fig. 5H**) in macrophages significantly compromised Mtb H37Rv burden (**Fig. 5G, H**). Furthermore, to identify the importance of each of the G9a-dependent cholesterol biosynthesis and uptake genes, each gene was individually knocked down and assessed for their effect on the expression of antioxidant genes (**Fig. S7**) as well as Mtb burden (**Fig. 5I**). Our observations suggest a dominant role for the genes *Hmgcs1* and *Aacs*, catalyzing the pioneer steps of cholesterol biosynthesis, in impacting oxidative stress and mycobacterial burden. These evidences highlight the critical functions of cholesterol accumulation in mycobacteria-infected hosts. However, further investigation into the exact implication of *Hmgcs1* and *Aacs* in regulating antioxidant gene expression is warranted.

**Figure 5.**
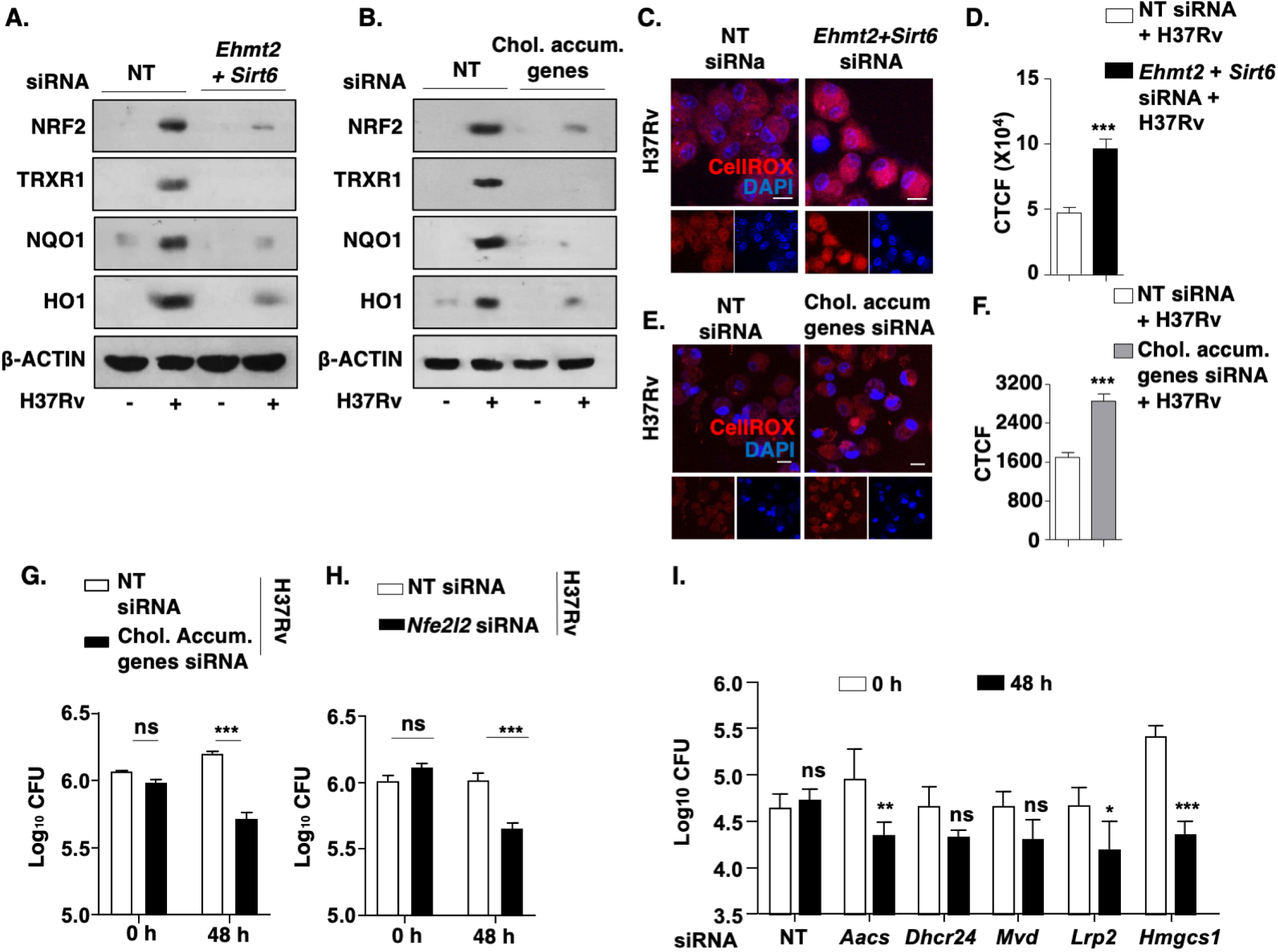
Cholesterol accumulation regulates the expression of anti-oxidant genes during mycobacterial infection. **(A-I)** BALB/c mouse peritoneal macrophages were transfected with NT or *Ehmt2* and *Sirt6* siRNA or *Nfe2l2* siRNA or cholesterol accumulation genes siRNA (*Lrp2*, *Aacs*, *Hmgcs1*, *Mvd* and *Dhcr24* siRNAs) followed by 48 h of H37Rv infection. **(A, B)** The expression of the indicated molecules was assessed at the protein level by immunoblotting. **(C-F)** CellROX staining was performed to assess ROS levels in macrophages; **(C, E)** representative images and **(D, F)** respective quantification (n=200-300). **(G, H, I)** *In vitro* CFU was assessed. The MOI of infection is 1:10 (macrophage: mycobacteria) for all the *in vitro* experiments. All data represent the mean ± SEM from 3 independent experiments. The blots are representative of 3 independent experiments. *, P < 0.05; **, P < 0.01; ***, P < 0.001; ns, not significant (Student’s t-test for C-E, G and I). Med, Medium; NT, non-targeting; chol. accum. genes, cholesterol accumulation genes. Scale bar, 10μm.

In the perspective of the above-mentioned observations, we explored the role for WNT signaling pathway during Mtb-driven expression of G9a and Sirt6. WNT pathway is known to modulate various cellular events like autophagy during Mtb infection^51^. Importantly, it has been associated with antioxidants such as NRF2 for defining neuronal developmental pathways^52^. Activated WNT signaling, driven by WNT3A, has also been shown to enhance NRF2-mediated anti-oxidant gene expression by preventing the GSK3β-dependent phosphorylation and subsequent proteasomal degradation of NRF2 in hepatocytes^53^. Further, its contribution in regulating lipid accumulation by endocytosis of LDL-derived cholesterol ^54^ indicated its possible role in yet another aspect of Mtb infection, i.e. cholesterol accumulation. G9a and Sirt6 expression was found to be dependent on Mtb H37Rv-activated WNT pathway (**Fig. S8A-C; S8A:** hallmarks of Mtb-activated WNT signaling: increased pGSK3β and reduced pβ-CATENIN); as inhibition of the pathway with pharmacological inhibitors (IWP2 and β-CATENIN inhibitor) (**Fig. S8C, right panel**) or knockdown of *Ctnbb1* (**Fig. S8C, middle panel; Fig. S8B:** knockdown validation) compromised the levels of G9a and Sirt6. Conversely, β-CATENIN over-expression alone induced the expression of the concerned histone modifiers (**Fig. S8C, left panel**). Further, β-CATENIN was found to be recruited to the promoters of *Ehmt2* and *Sirt6*, (**Fig. S8D**). siRNA-mediated knockdown of *Ctnbb1* compromised the ability of Mtb to differentially regulate cholesterol metabolism genes (**Fig. S8G**); subsequent cholesterol accumulation (**Fig. S8E, F**) and hence Mtb H37Rv survival (**Fig. S8H**). These findings indicate that Mtb infection leads to the WNT signaling pathway-dependent expression of G9a/Sirt6 as well as accumulation of cholesterol, which drives a secure niche for the pathogen to survive.

### G9a and Sirt6 contribute to mycobacterial pathogenesis

The observed G9a/Sirt6-dependent accumulation of cholesterol and the related abatement of mycobacterial burden upon their functional loss incited us to determine the impact of G9a and Sirt6 in defining *in vivo* Mtb burden and associated lung tissue pathology during Mtb H37Rv infection. We found that therapeutic treatment of Mtb H37Rv-infected mice with G9a inhibitor not only compromised cholesterol accumulation but also reduced mycobacterial CFU and led to a decreased level of Mtb infection-specific granulomatous lesions. Lung histopathological examination by Hematoxylin and Eosin (H and E) staining also revealed a marked reduction in the percentage of lung area covered with the characteristic TB granulomatous lesions, with an overall decline in total granuloma score compared to the untreated counterparts (**Fig. 6A-E**). Further, we observed limited Mtb H37Rv CFU in the lungs and spleen in *Sirt6* heterozygous mice and up to 50% restriction in the ability of *Sirt6* heterozygous mice to effectively develop TB granulomatous lesion (**Fig. 6F-J**). Therefore, the partial normalization of total lung architecture in mice lacking G9a or Sirt6 functions strongly indicates the relevance of the histone modifications conferred by G9a and Sirt6 in the pathogenesis of TB. We believe that thwarted cholesterol accumulation, leading to enhanced oxidative stress, jeopardizes mycobacterial survival strategies, thereby restricting overall TB progression in mice with abrogated G9a/Sirt6 functions.

**Figure 6.**
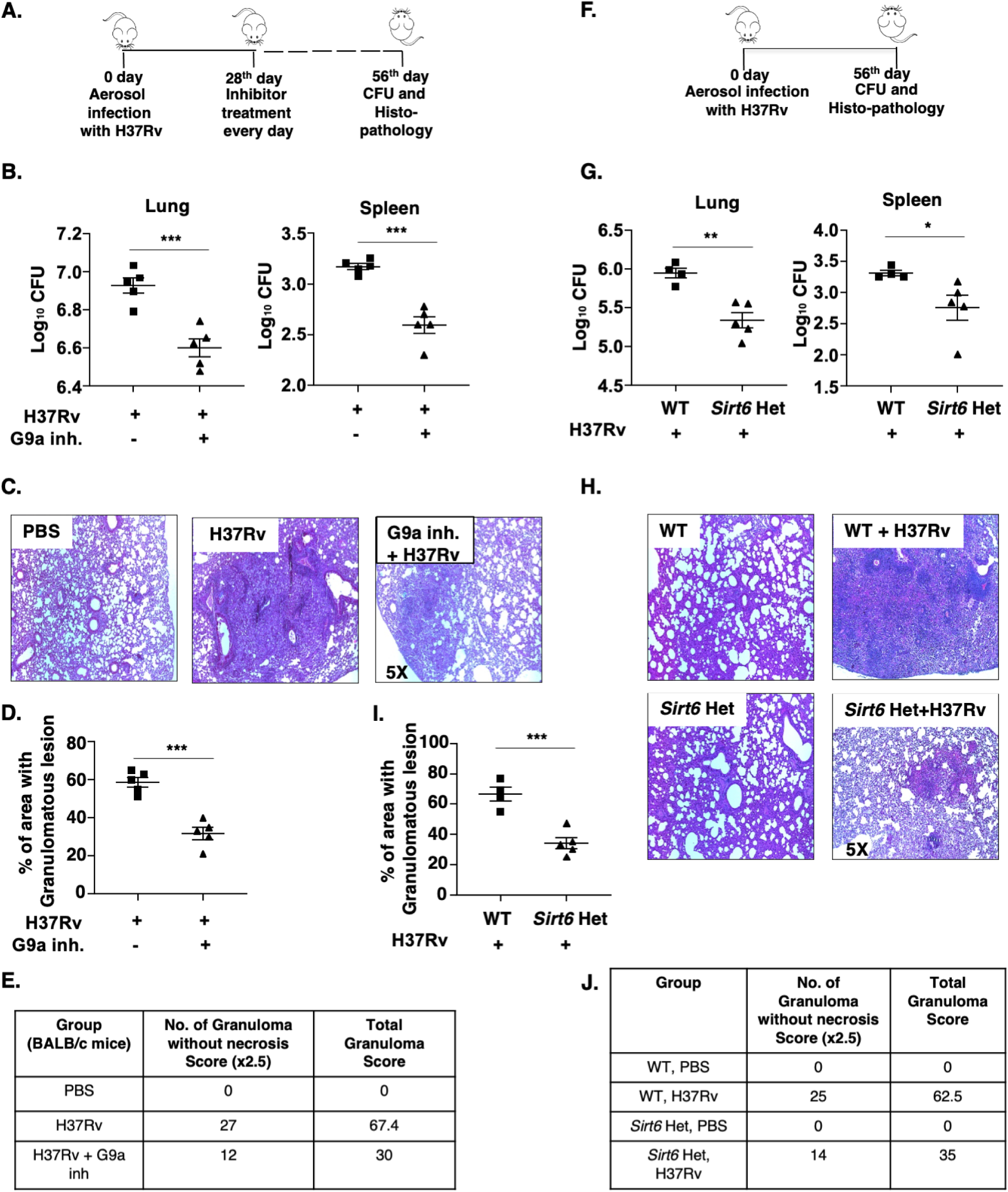
Epigenetic modifiers G9a and Sirt6 aid in mycobacterial pathogenesis. **(A-J)** Mice were aerosolized with 200 CFU of H37Rv. **(A)** Schematic of *in vivo* mouse TB model for G9a inhibitor therapeutic treatment. **(B)** CFU of H37Rv from G9a inhibitor (40mg/kg) treated and untreated BALB/c mice in lung and spleen, after 56 days of total infection and therapeutic treatment. **(C-D)** Lungs of BALB/c mice from the indicated groups were analyzed for TB pathology by H & E staining; **(C)** representative image, **(D)** % of granulomatous area and **(E)** corresponding histological evaluation for granuloma score. **(F)** Schematic of *in vivo* TB infection model for WT (littermate control) and *Sirt6* het mice. **(G)** CFU of H37Rv in lung and spleen of WT (littermate control) and *Sirt6* het mice after 56 days of H37Rv infection. Lungs from the indicated groups of mice were analyzed by H and E staining; **(H)** representative images, **(I)** % of granulomatous area and **(J)** corresponding histological evaluation for granuloma score. All data represents the mean ± SEM from 4-5 mice, *, P < 0.05; **, P < 0.01; ***, P < 0.001 (Student’s *t*-test for B, D, G and I). WT, Wild type; Het, heterozygous; inh., inhibitor.

## Discussion

The formation of FMs has been described as an integral part of TB pathogenesis and the constituents of the lipid droplets (LDs) contained in the FMs associate with diverse functions. Specifically, cholesterol uptake by Mtb and utilization to achieve survival advantages has been vividly elucidated^55–58^. We uncover the Mtb-driven host molecular players that lead to the accumulation of this essential factor in host cells during infection. Despite the presence of compelling evidences for the implication of cholesterol in the pathogenesis of Mtb, the epidemiological surveys depict a nonlinear and complex relationship between high cholesterol and TB progression^59^. Similarly, in mouse models of pulmonary TB, a cholesterol-rich diet (high serum cholesterol levels) has been related to distinct disease outcomes. For instance, in *ApoE* KO mice, high serum cholesterol impairs host defense against Mtb^60^; while that in *Ldlr* KO mice does not alter the capacity of the host to restrict mycobacterial replication^61^. These uncertainties may be explained by the differences in cholesterol availability that arise from its esterification or association with lipoproteins to form VLDLs, LDLs and HDLs. Therefore, a clear picture defining the role of cholesterol still warrants investigation.

The accumulation of cholesterol imparts regulatory effects on several aspects of host immunity by altering processes ranging from plasma membrane dynamics to maintaining serum cholesterol levels and epigenetic deregulations. Cholesterol is important for the adaptive immune system for its contribution to the formation of plasma membrane lipid rafts, which facilitate immune functions such as T-cell and B-cell signaling, their activation and proliferation^62,63^. Further, high serum cholesterol leads to autoimmune and inflammatory manifestations via aberrant immune activation^64^. Alongside these important roles, cholesterol accumulation also shapes the innate immune arm by modulating functions such as TLR signaling, monocyte proliferation, macrophage polarization, apoptosis as well as dendritic cell maturation and activation under distinct conditions^65–69^, including infections. For instance, cholesterol has been shown to play a crucial role in regulating Salmonella-induced autophagy ^70^ and lowering free cholesterol by their conversion to oxysterols has been implicated in providing immunity against *Listeria monocytogenes* infection^71^.

During mycobacterial infection, in particular, suppression of intracellular cholesterol accumulation via oxysterols (natural LXR activators) or by inhibition of SREBP2 has been shown to enhance the production of anti-microbial peptides and restrict Mtb burden^72^. Inline, loss of function of LXRα and LXRβ (leading to reduced expression of *Abca1*) has been reported to render mice more susceptible to Mtb infection due to defective recruitment of innate effector cells and innate immune functions as well as severely compromised Th1/Th17 functions^73^. In the current study, we present a novel mechanism by which free cholesterol accumulated within host cells can aid in

Mtb pathogenesis. We show that G9a-SREBP2 and SIRT6 independently regulates the cholesterol homeostasis wherein G9a-SREBP2 regulates cholesterol biosynthesis genes, whereas, SIRT6 regulates the expression of cholesterol efflux genes but has no effect on cholesterol biosynthesis genes. We find that cholesterol accumulation modulates the innate immune arm by driving the expression of anti-oxidative genes that would favor Mtb survival by circumventing oxidative stress responses and mediators. This aligns with the observation that cells with high cholesterol upregulate anti-oxidants such as NRF2 and HO-1 to mitigate oxidative stress^74^. A recent report from our lab proposes that mycobacterial clearance pathways such as apoptosis and pro-inflammatory cytokine production are hampered by classical anti-oxidative molecules TRXR1 and NQO1^5^. Therefore, cholesterol-dependent antioxidant production and subsequent innate and adaptive immune alterations not reported as yet, can potentially help in strengthening the understanding of the survival strategies employed by Mtb.

In the light of host-directed therapeutics, our finding is in congruence with a previous study where statins, that decrease cholesterol levels by inhibiting HMGCoA reductase (a rate-limiting step of cholesterol biosynthesis), had been reported to inhibit mycobacterial growth^75^. With the individual knockdown of G9a-dependent cholesterol biosynthesis genes, we tease out the specific contribution of *Hmgcs1* and *Aacs* in regulating cholesterol-driven mitigation of oxidative stress and subsequently, mycobacterial burden. Therefore, this study provides an avenue for testing alternate targets for effective combinatorial therapy against TB and for dedicated studies on metabolic homeostasis and mycobacterial pathogenesis in *Hmgcs1* or *Aacs* knockout conditions. Recently, mammalian sirtuins have been proposed as a potential target for host-directed therapy against TB. For example, SIRT1 activators ameliorates lung pathology, SIRT3 promotes antimycobacterial responses whereas SIRT2 inhibition has been shown to reduce Mtb burden ^25,76,77^. In the current study, we find that SIRT6 benefits the Mtb survival and impacts lung pathology, thereby establishing the class of sirtuins as potential targets for TB therapeutics.

Together, we report that epigenetic modifiers G9a and Sirt6 are induced by Mtb, and the two enzymes differentially occupy the promoters of distinct arms of cholesterol biosynthesis, uptake and efflux genes, in order to build up cholesterol within host cells. Interception of G9a and Sirt6 restricts mycobacterial burden and limits TB pathology, plausibly by compromising free cholesterol accumulation and thereby increasing oxidative stress in host cells (**Fig. 7**). We believe that an organ-specific and carefully titrated delivery of therapeutics against these epigenetic factors would provide rational and clinically relevant adjuvants for TB treatment.

**Figure 7.**
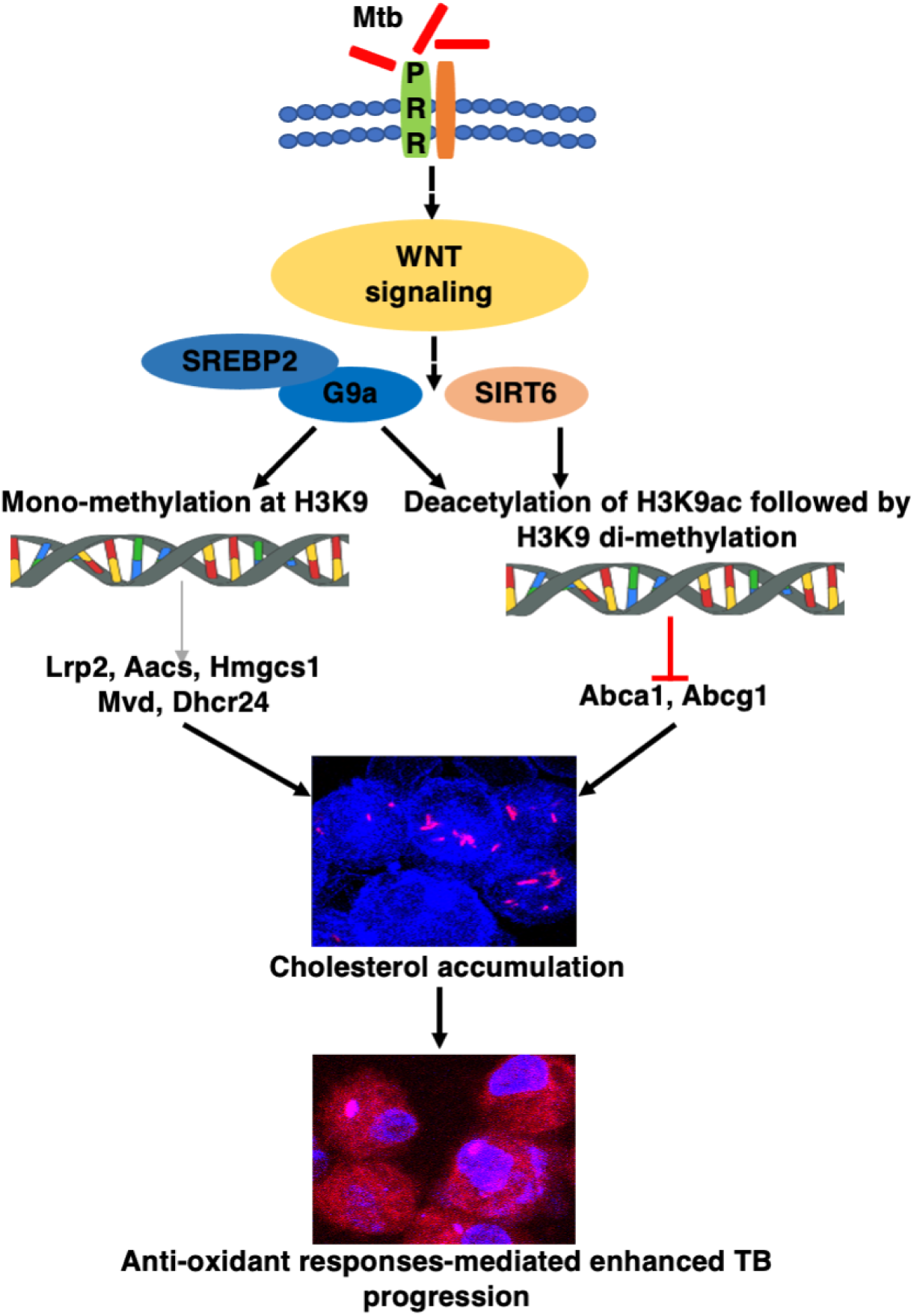
Schematic, Mycobacteria utilizes the host epigenetic factors G9a and SIRT6 to augment cholesterol accumulation and antioxidant responses in order to aid its survival within the host.

## Materials and Methods

### Mice and Cells

Male and female mice of the following strains were utilized in the study: BALB/c (stock number 000651, The Jackson Laboratory, USA), *Sirt6* KO (kind gift from Dr. Ullas Kolthur-Seetharam, TIFR, India and Dr. Nagalingam Ravi Sundaresan, IISc, India; primary source: The Jackson Laboratory, USA, stock number 006050) and *Sirt6* heterozygous (kind gift from Dr. Ullas Kolthur-Seetharam, TIFR, India and Dr. Nagalingam Ravi Sundaresan, IISc, India; primarily generated by crossing *Sirt6* KO mice with WT 129S6 mice). Mouse primary macrophages were isolated from peritoneal exudates using ice-cold PBS four days post intraperitoneal injection of 1.5ml of brewer thioglycollate (8%). RAW 264.7 mouse macrophages cell line was obtained from National Center for Cell Sciences, Pune, India; and used for transient transfection experiments using plasmids as they are better suited for transfection as compared to the peritoneal macrophages that are known to be highly sensitive to external DNA^78,79^. Primary macrophage and RAW 264.7 macrophage cell line was cultured in Dulbecco’s Minimal Eagle Medium (Gibco, Thermo Fisher Scientific) supplemented with 10% heat-inactivated Fetal Bovine Serum (Gibco, Thermo Fisher Scientific) and maintained at 37°C in 5% CO_2_ incubator. All strains of mice were obtained from The Jackson Laboratory and maintained in the Central Animal Facility (CAF), Indian Institute of Science (IISc) under 12 h light and dark cycle.

### Ethics Statement

Experiments involving mice and virulent mycobacteria (Mtb H37Rv) were carried out after the approval from Institutional Ethics Committee for animal experimentation and Institutional Biosafety Committee. The animal care and use protocol adhered were approved by national guidelines of the Committee for the Purpose of Control and Supervision of Experiments on Animals (CPCSEA), Government of India.

### Bacteria

Mtb H37Rv was a kind research gift from Prof. Kanury Venkata Subba Rao, THSTI, India. tdTomato Mtb H37Rv was a kind research gift from Dr. Amit Singh, IISc, India. Mycobacterial cultures were grown to mid-log phase in Middlebrook 7H9 medium (Difco, USA) supplemented with 10% OADC (oleic acid, albumin, dextrose, catalase) and hygromycin for tdTomato Mtb H37Rv. Single-cell suspensions of mycobacteria were obtained by passing mid-log phase culture through 23, 28 and 30 gauge needle 10 times each and used at a multiplicity of infection 10 unless mentioned otherwise. The studies involving virulent mycobacterial strains were carried out at the biosafety level 3 (BSL-3) facility at CIDR, IISc.

### Reagents and antibodies

All general laboratory chemicals were obtained from Sigma Aldrich/Merck Millipore, Thermo Fisher Scientific, HiMedia or Promega. Tissue culture plasticware was purchased from Jet Biofil or Tarsons Products Pvt Ltd. Further details are provided in the supplementary file.

### Transient transfection studies

RAW 264.7 macrophages were transiently transfected with 5μg of overexpression constructs of β-CATENIN and SIRT6; or peritoneal macrophages were transfected with 100 nM each of siGLO Lamin A/C, non-targeting siRNA or specific siRNAs against *Ehmt2*, *Sirt6*, *Ctnnb1*, *Lrp2*, *Aacs, Hmgcs1, Mvd, Dhcr24*, *Srebf2*, *Nfe2l2* (purchased from Dharmacon as siGENOMETM SMARTpool reagents) with polyethyleneimine. 70-80% transfection efficiency was observed by counting the number of siGLO Lamin A/C positive cells in a microscopic field using fluorescence microscopy. 36 h post-transfection (for experiments with RAW 264.7 cells) or 24h post-transfection (for experiments with peritoneal macrophages), the cells were treated or infected as indicated and processed for analyses.

### *In vivo* mouse model and inhibitor treatment

BALB/c mice (n=40) were infected with mid-log phase Mtb H37Rv, using a Madison chamber aerosol generation instrument calibrated to 200 CFU/animal. Aerosolized animals were maintained in a securely commissioned BSL3 facility. Post 28 days of established infection, mice were administered a daily dose of G9a inhibitor BIX-01294 (40mg/kg) ^80^ intra-peritoneally for 28 days. Alternately, wild type (littermate control) mice or *Sirt6* heterozygous mice were infected as described above. In each case, on the 56^th^ day, mice were sacrificed, spleen and left lung lobe and spleen were homogenized in sterile PBS, serially diluted and plated on 7H11 agar containing OADC to quantify CFU. Upper right lung lobes were fixed in formalin, embedded in paraffin and stained with hematoxylin and eosin and immunofluorescence analysis. For Granuloma scoring, different scores were assigned based on the characteristic granulomatous features that is granuloma with necrosis (Score=5), without necrosis (Score = 2.5) and with fibrosis (Score = 1)^81^. For total granuloma scoring, the number of granulomas in each lung lobe was multiplied with the characterized feature score. The granulomatous area of lung sections stained with H&E was measured using Image J software (granulomatous area/ total area *100).

### Statistical analysis

Levels of significance for comparison between samples were determined by the Student’s t-test and one-way ANOVA followed by Tukey’s multiple-comparisons. The data in the graphs are expressed as the mean ± SEM for the values from at least 3 or more independent experiments and P values < 0.05 were defined as significant. GraphPad Prism 6.0 software (GraphPad Software, USA) was used for all the statistical analyses.

**All the details concerning the pharmacological reagents, antibodies, *in vitro* experiments and procedures have been provided as supplementary information**

## Acknowledgments

We thank CAF, IISc for providing mice for experimentation. β-CATENIN cDNA was gifted by Dr. Roel Nusse, Stanford University School of Medicine, USA. We acknowledge the BSL-3 facility for allowing the experiments on Mtb H37Rv to be carried out.

## Abbreviations

HMT: histone methyl transferase
TB: tuberculosis
FM: foamy macrophage
ER: endoplasmic reticulum
HDAC: histone deacetylase
CFU: colony forming unit
BMDM: bone marrow derived macrophage
KO: knock out
LRP: Low density lipoprotein receptor-related protein 2
SREBP: Sterol response element binding protein
ABC: ATP-binding cassette
LDL: low density lipoprotein
LD: lipid droplet
HDL: high density lipoprotein
NRF2: nuclear factor erythroid 2 (NFE2) related factor 2
VLDL: very low density lipoprotein
PBMC: polymorphic blood mononuclear cells

## SUPPLEMENTARY FIGURES AND LEGENDS

**Figure S1.**
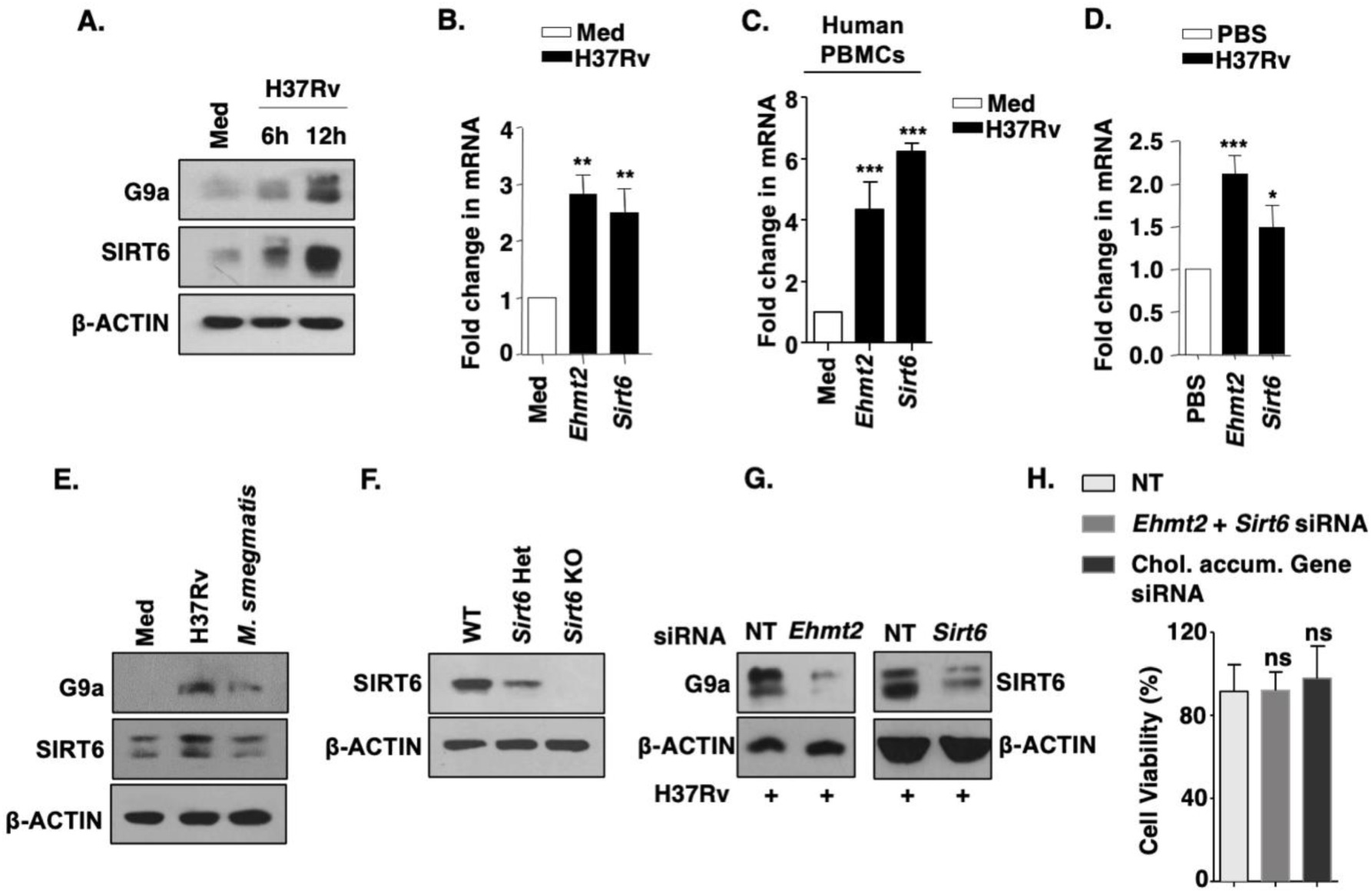
Mtb-triggered expression of epigenetic modifiers G9a and Sirt6 in host cells. **(A)** Immunoblot of G9a and SIRT6 in BALB/c mouse peritoneal macrophages infected with H37Rv for 6 h or 12 h. **(B, D)** Transcript level of the *Ehmt2* and *Sirt6* was analyzed by qRT-PCR **(B)** in BALB/c mouse peritoneal macrophages infected with H37Rv for 12 h or **(C)** in human PBMCs infected with H37Rv for 12 h or **(D)** in lung homogenates of mice infected with H37Rv for 56 days. **(E)** Protein level of G9a and SIRT6 was assessed in BALB/c macrophages infected with H37Rv or *M. smegmatis* for 12 h by immunoblotting. **(F)** The protein levels of SIRT6 was assessed in lung homogenates of WT (littermate control), *Sirt6* het and *Sirt6* KO mice by immunoblotting. **(G)** BALB/c mouse peritoneal macrophages were transfected with the indicated siRNAs and infected with H37Rv for 12 h. Whole cell lysates were assessed for the knock down of G9a and SIRT6 by immunoblotting. **(H)** MTT assay was performed to assess cell viability of BALB/c macrophages transfected with NT or *Ehmt2* and *Sirt6* siRNA or chol. accum. genes siRNA (*Lrp2*, *Aacs*, *Hmgcs1*, *Mvd* and *Dhcr24*). The MOI of infection is 1:10 (macrophage: mycobacteria) for all the *in vitro* experiments. All data represents the mean ± SEM from 3 independent experiments. The blots are representative of 3 independent experiments. *, P < 0.05; **, P < 0.01; ***, P < 0.001 (Student’s t-test for B, C, D and H). Med, Medium. NT, non-targeting; ns, not significant; WT, wild type; Het, heterozygous; KO, knock out; chol. accum. gene, cholesterol accumulation genes.

**Figure S2.**
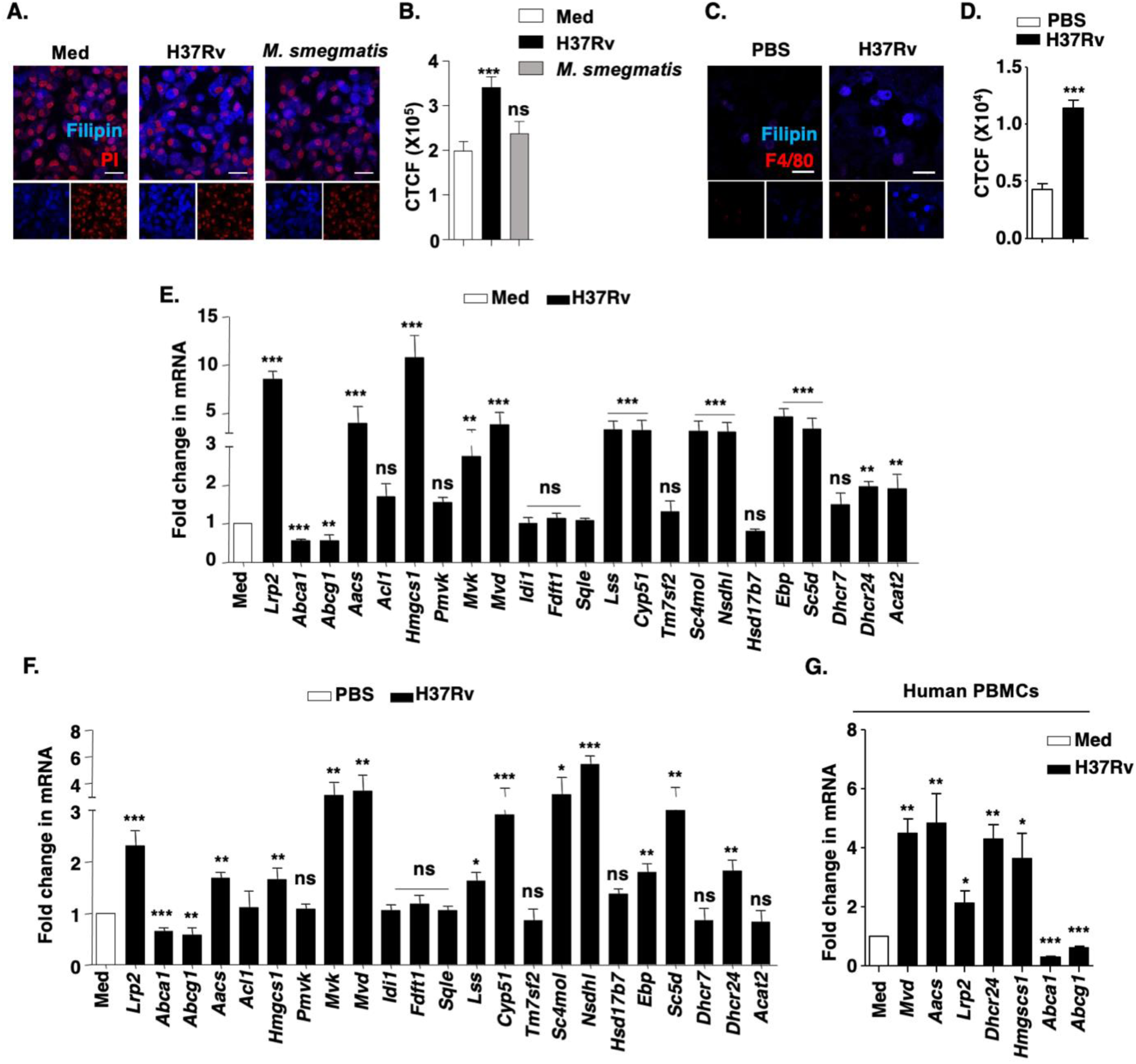
Mtb-driven free cholesterol accumulation in host cells. **(A, B)** BALB/c mouse peritoneal macrophages were infected with H37Rv or *M. smegmatis* for 48 h and assessed for cholesterol accumulation by Filipin staining; **(A)** representative image and **(B)** respective quantification (n=200-300). **(C)** Lung cryosections from BALB/c mice infected with H37Rv for 56 days was assessed for cholesterol accumulation by Filipin staining in macrophages stained with F4/80, **(D)** quantification of Filipin staining in F4/80 positive cells in lung cryosections. **(E, F)** Transcript level of the indicated set of genes was analysed by qRT-PCR **(E)** in BALB/c mouse peritoneal macrophages infected with H37Rv for 12 h, **(F)** in lung homogenates of BALB/c mice infected with H37Rv for 56 days and **(G)** in human PBMCs infected with H37Rv for 12 h. The MOI of infection is 1:10 (macrophage:mycobacteria) for all the *in vitro* experiments. All data represents the mean ± SEM from 3 independent experiments. Confocal images were obtained from lungs of at least three groups of mice. *, P < 0.05; **, P < 0.01; ***, P < 0.001 (One-way ANOVA for B, Student’s t-test for D-F). Med, Medium; CTCF; corrected total cell fluorescence; MFI, mean fluorescence intensity; PI, Propidium Iodide (nuclear stain); PBMC, peripheral blood mononuclear cells; Scale bar, 25 μm.

**Figure S3.**
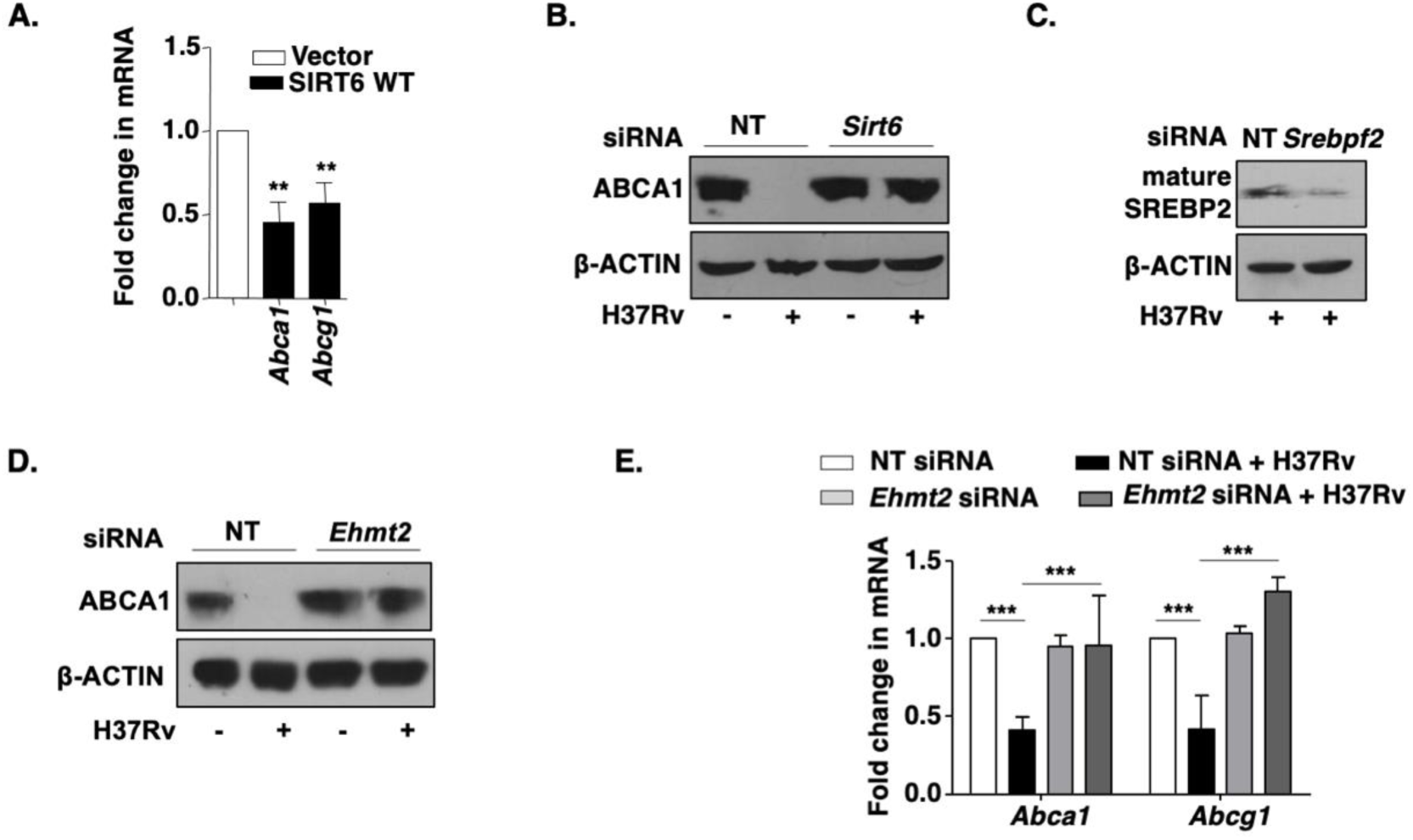
Interplay of G9a and Sirt6 in regulation of cholesterol efflux during Mtb infection. **(A)** RAW 264.7 macrophages were transfected with vector or SIRT6 WT construct and transcript levels of ABC transporters was analysed by qRT-PCR. **(B-E)** BALB/c mouse peritoneal macrophages were transfected with NT or *Ehmt2* or *Sirt6* or *Srebf2* siRNA as indicated, followed by 12 h infection with H37Rv. Whole cell lysates were assessed for **(B)** ABCA1 or **(C)** SREBP2 expression by immunoblotting. **(D)** ABCA1 was assessed by western blotting and **(E)** transcript level of the indicated genes were measured by qRT-PCR. The MOI of infection is 1:10 (macrophage: mycobacteria) for all the *in vitro* experiments. All data represents the mean ± SEM from 3 independent experiments. The blots are representative of 3 independent experiments. **, P < 0.01; ***, P < 0.001 (Student’s t-test for A, One-way ANOVA for D and E); Med, Medium; NT, non-targeting; ns, not significant.

**Figure S4.**
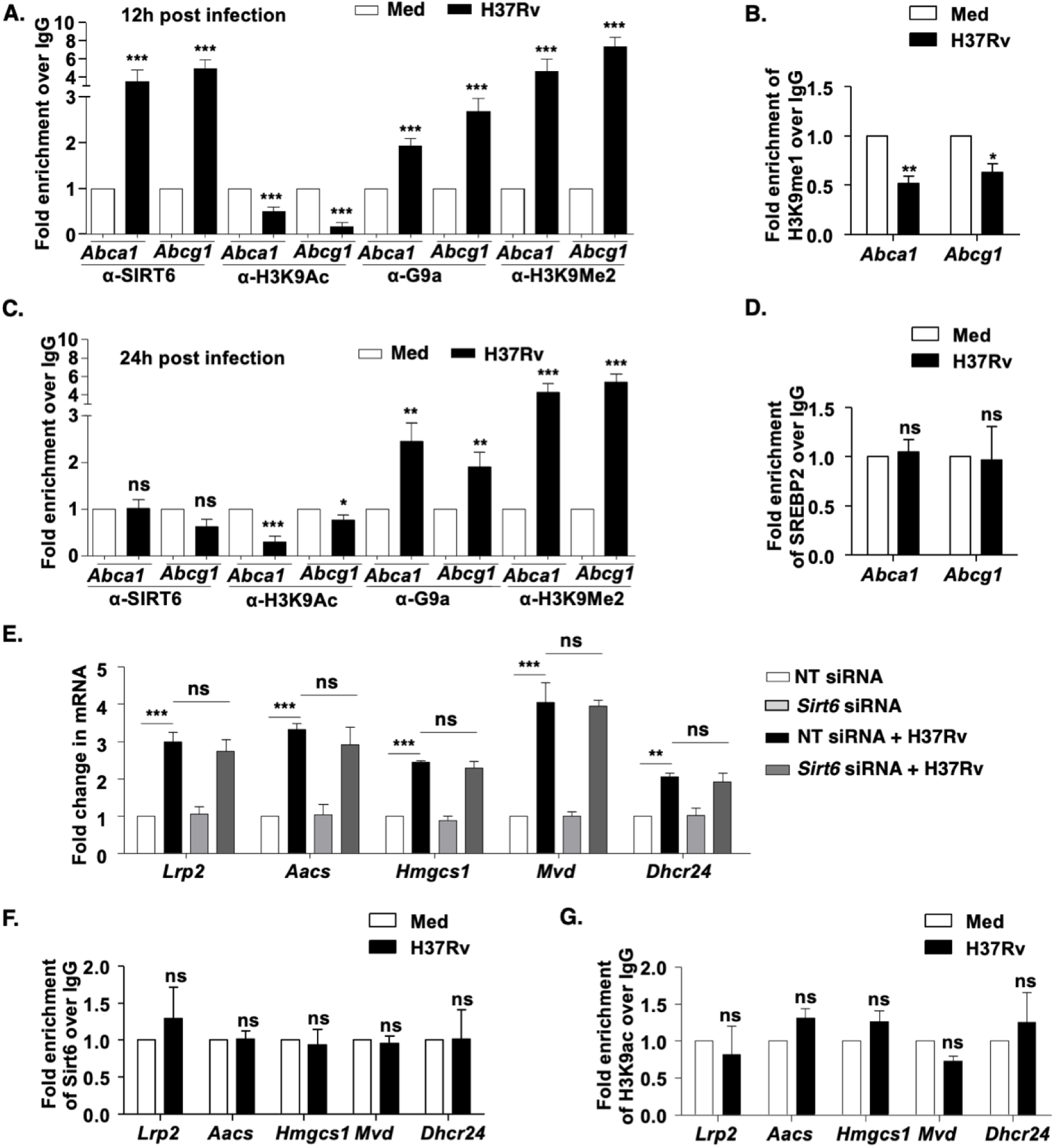
G9a and Sirt6 mediated transcriptional regulation of cholesterol efflux and synthesis genes. **(A, B)** BALB/c mouse peritoneal macrophages were infected with H37Rv for 12 h, and assessed for the recruitment of SIRT6, G9a and presence of H3K9Ac H3K9me2 and H3K9me1 **(B)**, on the promoters of *Abca1* and *Abcg1*. **(C)** BALB/c mouse peritoneal macrophages were infected with H37Rv for 24h and assessed for recruitment of SIRT6, G9a, H3K9Ac and H3K9me2 on promoters of *Abca1* and *Abcg1* **(D)** ChIP analysis of SREBP2 on the promoters of *Abca1* and *Abcg1* in BALB/c mouse peritoneal macrophages after 12 h H37Rv infection. **(E)** Cholesterol accumulation genes were assessed in BALB/c mouse peritoneal macrophages transfected with NT or *Sirt6* siRNA post 12h of H37Rv infection. **(F, G)** Recruitment of SIRT6 and H3K9Ac on the promoters of cholesterol accumulation genes in BALB/c mouse peritoneal macrophages upon 12 h of H37Rv infection. The MOI of infection is 1:10 (macrophage: mycobacteria) for all the *in vitro* experiments. All data represents the mean ± SEM from 3 independent experiments, *, P < 0.05; **, P < 0.01; ***, P < 0.001 (Student’s t-test in A-F). Med, Medium; NT, non-targeting; ns, not significant.

**Figure S5.**
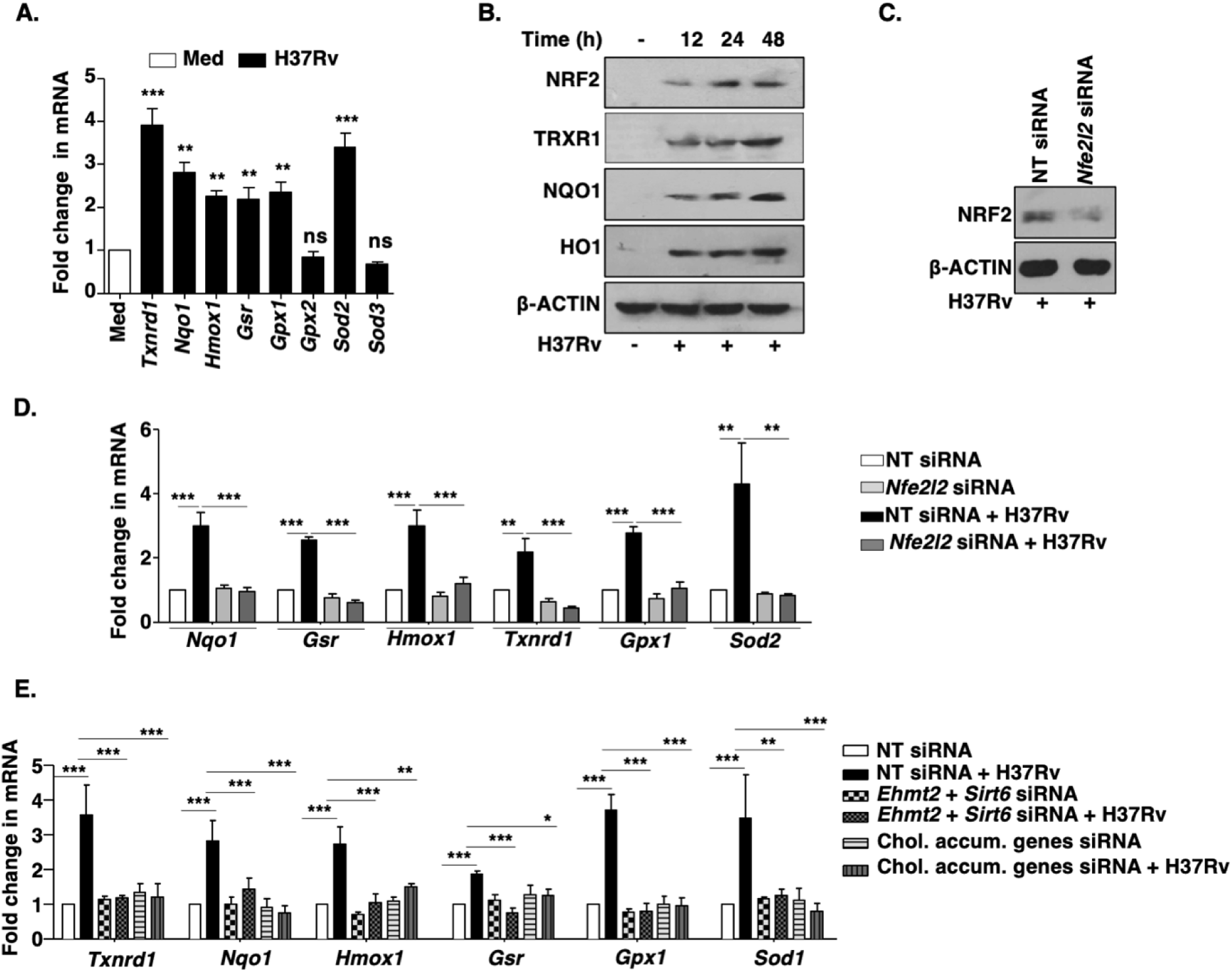
NRF2 and its target genes are expressed during Mtb infection. **(A)** BALB/c mouse peritoneal macrophages were infected with H37Rv for 48 h and the expression of NRF2 target genes was assessed by qRT-PCR. **(B)** BALB/c mouse peritoneal macrophages were infected with H37Rv for the indicated time points and whole cell lysates were assessed for the expression of NRF2 and its target genes. **(C)** Immunoblotting to validate NRF2 knockdown in murine macrophages transfected with *Nfe2l2* siRNA. **(D-E)** BALB/c mouse peritoneal macrophages were transfected with NT or **(D)** *Nfe2l2* siRNA or **(E)** *Ehmt2* and *Sirt6* siRNA or Chol accum genes siRNA (combination of *Lrp2*, *Aacs*, *Hmgcs1*, *Mvd* and *Dhcr24* siRNAs) in the presence or absence of 48 h infection with H37Rv and the transcript levels of NRF2 target genes were analysed by qRT-PCR. The MOI of infection is 1:10 (macrophage:mycobacteria) for all the *in vitro* experiments. All data represents the mean ± SEM from 3 independent experiments; *, P < 0.05; **, P < 0.01; ***, P < 0.001 (Student’s t-test for A and One-way ANOVA for D, E) and the blots are representative of 3 independent experiments. Med, Medium; NT, non-targeting; ns, not significant; chol. accum. genes, cholesterol accumulation genes.

**Figure S6.**
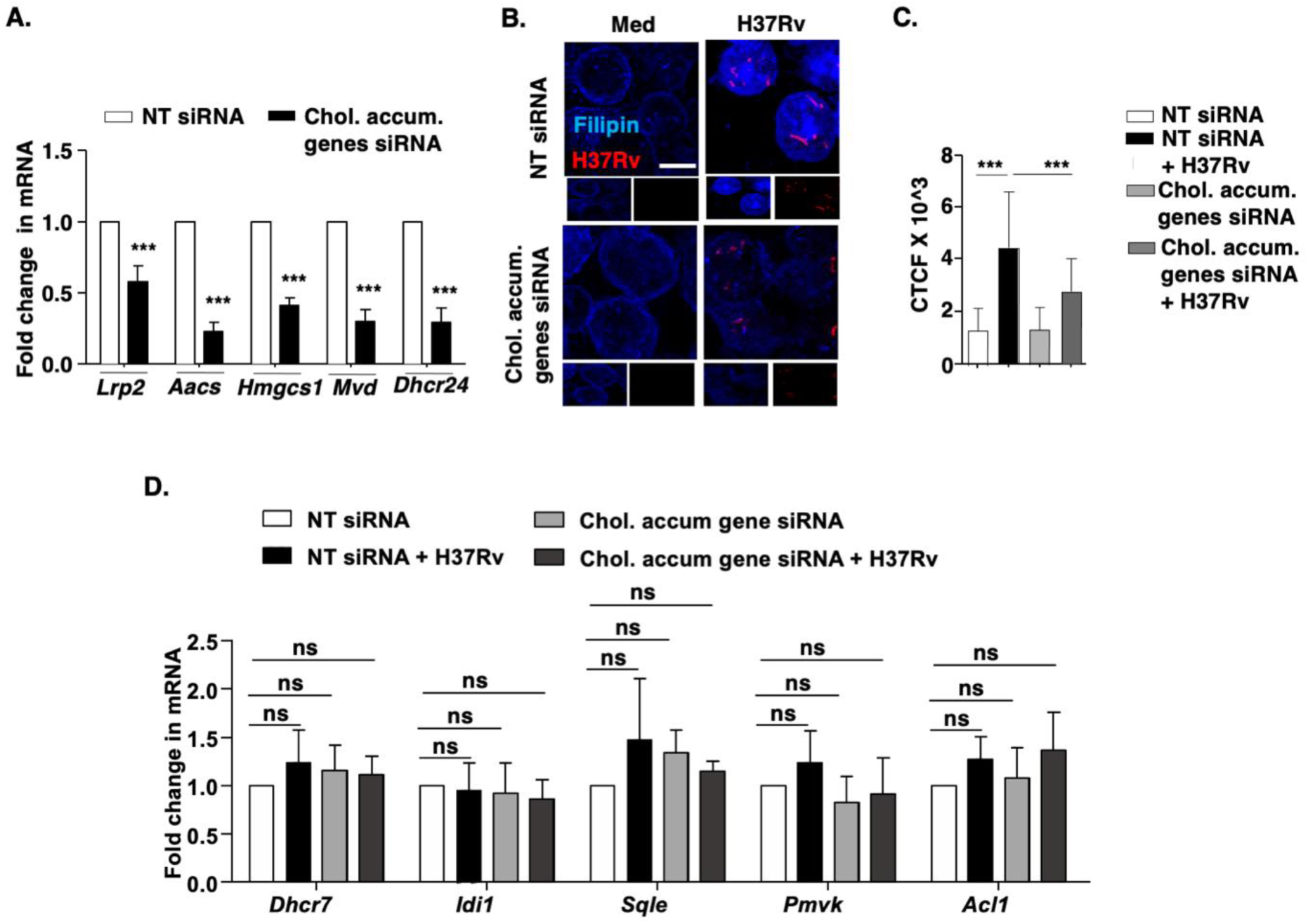
Validation of abrogated cholesterol accumulation in cells knocked down for genes involved in cholesterol biosynthesis and uptake. **(A-D)** BALB/c mouse peritoneal macrophages were transfected with siRNAs against the selected cholesterol accumulation genes (combination of *Lrp2*, *Aacs*, *Hmgcs1*, *Mvd* and *Dhcr24* siRNAs) or NT and **(A)** the expression of the concerned cholesterol genes was analysed by qRT-PCR; **(B-D)** followed by H37Rv infection for 48 h and cholesterol accumulation was confirmed by Filipin staining; **(B)** representative image, **(C)** respective quantification (n=200-300); **(D)** qRT-PCR to assess transcript levels of indicated genes. The MOI of infection is 1:10 (macrophage:mycobacteria) for all the *in vitro* experiments. All data represents the mean ± SEM from 3 independent experiments, ***, P < 0.001 (Student’s t-test for A; and One-way ANOVA for C); CTCF, corrected total cell fluorescence; NT, non-targeting; ns, not significant; chol. accum. gene, cholesterol accumulation genes. Scale bar, 10 μm.

**Figure S7.**
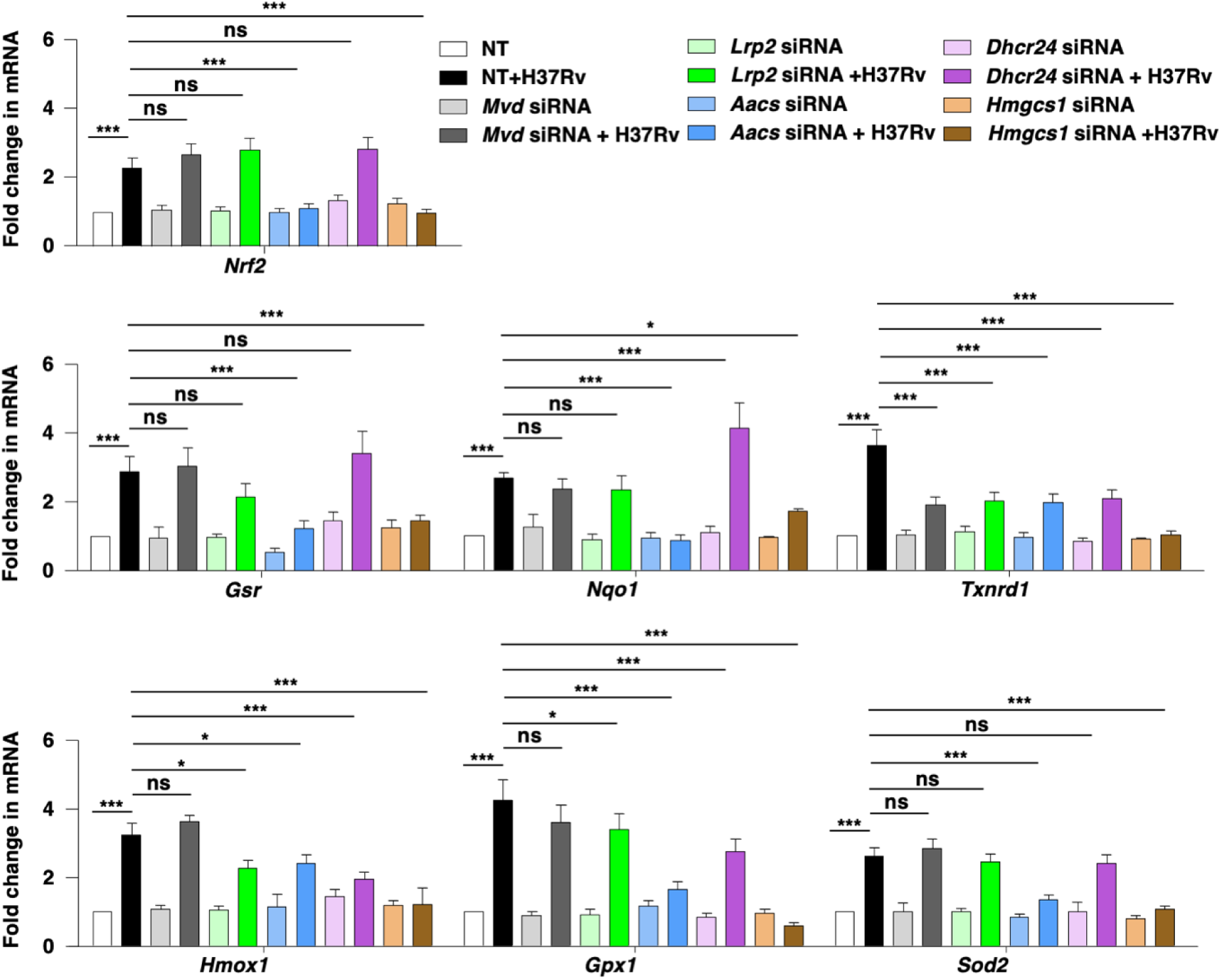
Cholesterol biosynthesis/uptake pathway modulates NRF2 and its target genes. BALB/c mouse peritoneal macrophages were transfected with NT or *Lrp2* or *Aacs* or *Hmgcs1* or *Mvd* or *Dhcr24* siRNAs and the transcript level of NRF2 and its target genes were assessed 48 h post Mtb infection. The MOI of infection is 1:10 (macrophage: mycobacteria) for all the *in vitro* experiments. Data represents the mean ± SEM from 3 independent experiments; *, P < 0.05; ***, P < 0.001 (Two-way ANOVA); NT, non-targeting; ns, not significant.

**Figure S8:**
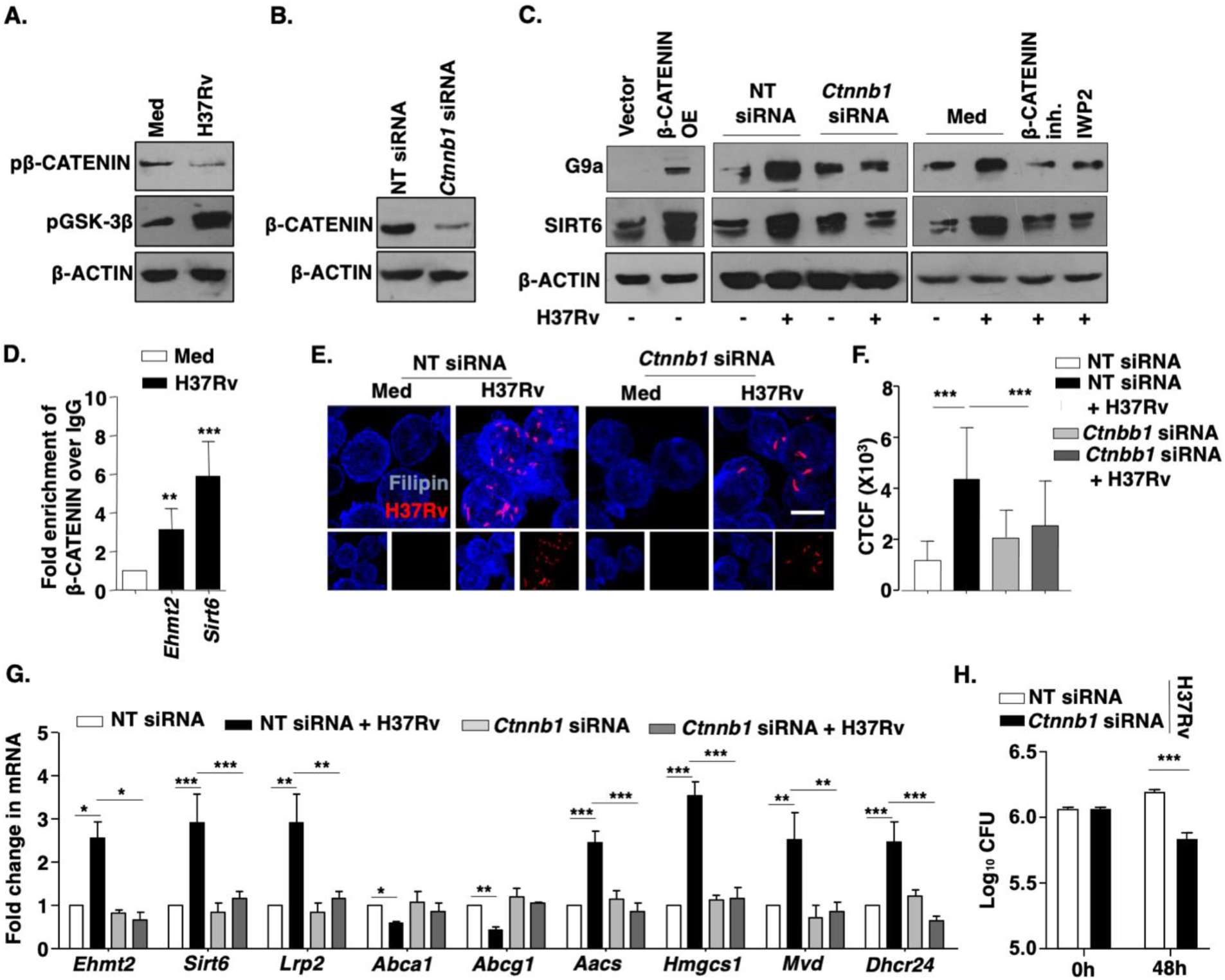
Contribution of WNT/β-CATENIN axis in mycobacterial infection. **(A)** BALB/c mouse peritoneal macrophages were infected with H37Rv for 1 h and whole cell lysates were assessed for the activation of WNT pathway. **(B)** Immunoblotting to validate β-CATENIN knockdown in BALB/c mouse macrophages transfected with NT or *Ctnnb1* siRNA. **(C)** RAW 264.7 macrophages were transfected with β-CATENIN OE construct (**C**, left panel) or mouse peritoneal macrophages were transfected with NT or *Ctnnb1* siRNA (**C**, middle panel) or BALB/c mouse peritoneal macrophages were pre-treated with β-CATENIN inhibitor (15 μM) or IWP2 (5 μM) for 1 h (**C**, right panel), followed by 12 h infection with H37Rv. Whole cell lysates were assessed for SIRT6 and G9a expression by immunoblotting. **(D)** β-CATENIN recruitment at the promoter of *Ehmt2* and *Sirt6* was assessed by ChIP assay in BALB/c mouse primary macrophages infected with H37Rv for 12 h. **(E, F)** Free cholesterol was assessed by Filipin staining in BALB/c mouse peritoneal macrophages transfected with NT or *Ctnnb1* siRNA followed by 48 h infection with tdTomato H37Rv, **(E)** representative image and **(F)** respective quantification (n=200-300). **(G)** Indicated genes were analyzed at transcript level by qRT-PCR in BALB/c mouse peritoneal macrophages that were transfected with NT or *Ctnnb1* siRNA followed by infection with H37Rv for 12 h. **(H)** BALB/c mouse macrophages were transfected with NT or *Ctnnb1* siRNA and *in vitro* CFU was assessed at the indicated time points post H37Rv infection. The MOI of infection is 1:10 (macrophage:mycobacteria) for all the *in vitro* experiments. All data represents the mean ± SEM from 3 independent experiments, *, P < 0.05; **, P < 0.01; ***, P < 0.001 (Student’s t-test for D; One-way ANOVA for F-H). All blots are representative of 3 independent experiments. Med, medium; β-CATENIN OE, β-CATENIN over expression; NT, non-targeting; inh., inhibitor. Scale bar, 10 μm.

## MATERIALS AND METHODS

### Antibodies

HRP-conjugated anti-β-ACTIN antibody, Filipin and 4′,6-Diamidino-2-phenylindole dihydrochloride (DAPI) were purchased from Sigma-Aldrich/ Merck Millipore. Alexa488-conjugated anti-rabbit IgG, HRP-conjugated anti-rabbit total IgG and light chain specific IgG antibodies were purchased from Jackson ImmunoResearch, USA; PE-conjugated F4/80 was procured from Tonbo Biosciences, USA. Anti-G9a, anti-SIRT6, anti-H3K9me1, anti-H3K9me2, anti-H3K9Ac, anti-Ser33/37/Thr41 phospho-β-CATENIN, anti-Ser9 phospho-GSK-3β, anti-β-CATENIN, anti-NRF2, anti-HO1 and anti-TRXR1 antibodies were obtained from Cell Signaling Technology, USA. Anti-LRP2 antibody was purchased from Santa Cruz Biotechnology, USA; anti-SREBP2 antibody was procured from Abcam, USA; and anti-NQO1 antibody was purchased from Calbiochem, USA.

### Treatment with pharmacological reagents

Cells were treated with concerned pharmacological inhibitors for 1 h prior to the experiment at the following final concentrations: BIX-01294 (G9a inhibitor, 5 μM), β-CATENIN inhibitor (15 μM), IWP-2 (5 μM). DMSO at 0.1% concentration was used as the vehicle control. In all experiments involving pharmacological reagents, a tested concentration was used after careful titration experiments assessing the viability of the macrophages using the MTT (3-(4,5-Dimethylthiazol-2-yl)-2,5-diphenyltetrazolium bromide) assay.

### MTT assay

siRNA transfected mouse peritoneal macrophages were treated with 3-(4, 5-Dimethylthiazol-2-yl)-2, 5-diphenyltetrazolium bromide (MTT) for 4 h at a final concentration of 0.5 mg/ml. Media was gently removed post incubation and 200 µL of DMSO was added. This solubilized purple formazan crystals were quantified by measuring absorbance at 550 nm in an 96-well plate reader. Viability of siRNA transfected macrophages were assessed relative to non-transfected macrophages.

### Isolation of Human PBMCs

Histopaque-1077 (Sigma-Aldrich, USA) polysucrose solution was utilized to isolate PBMCs from whole blood as per manufacturer’s instruction. Briefly, 3 ml of whole blood was carefully layered onto 3 ml of Histopaque-1077 in a 15 ml conical centrifuge tube followed by centrifugation at 400 × g for 30 min at room temperature. Upper layer was carefully removed without disturbing the opaque interface of mononuclear cells. The interface was transferred into a fresh 15 ml conical centrifuge tube and resuspended in 10 ml isotonic phosphate buffered saline solution. The solution was centrifuged at 250 × g for 10 min, cell pellet was resuspended and cultured in RPMI supplemented with 10 % heat inactivated FBS (Gibco-Life Technologies) in the presence of 10 ng/ml M-CSF (PeproTech, USA) for 5 days at 37 °C in 5 % CO_2_ incubator and utilized for experiments.

### RNA isolation and Real-Time qRT-PCR

Total RNA from treated, untreated and infected macrophages were isolated using TRI reagent (Sigma). 2 µg of total RNA was converted into cDNA using First Strand cDNA synthesis kit (Applied Biological Materials Inc.). Target gene expression was assessed by Real-Time quantitative Reverse Transcription-PCR (qRT-PCR) using SYBR Green PCR mix (Thermo Fisher Scientific). All the experiments were repeated at least 3 times independently to ensure the reproducibility of the results. *Gapdh* was used as internal control. The list of primers is detailed in Supplementary Tables 1 and 2.

### Immunoblotting analysis

Cells post treatment and/or infection were washed with 1X PBS. Whole cell lysate was prepared by lysing in RIPA buffer [50 mM Tris-HCl (pH 7.4), 1 % NP-40, 0.25 % sodium deoxycholate, 150 mM NaCl, 1 mM EDTA, 1 mM PMSF, 1 μg/ml each of aprotinin, leupeptin, pepstatin, 1 mM Na_3_VO_4_, 1 mM NaF] on ice for 30 min. Total protein from whole cell lysates was estimated by Bradford reagent. Equal amount of protein from each cell lysate was resolved on 12 % SDS-PAGE and transferred onto PVDF membranes (Millipore) by semi-dry immunoblotting method (Bio-Rad). 5 % non-fat dry milk powder in TBST [20 mM Tris-HCl (pH 7.4), 137 mM NaCl, and 0.1 % Tween 20] was used for blocking nonspecific binding for 60 min. After washing with TBST, the blots were incubated overnight at 4 °C with primary antibody diluted in TBST with 5 % BSA. After washing with TBST, blots were incubated with secondary antibody conjugated to HRP for 4 h at 4 °C. The immunoblots were developed with enhanced chemiluminescence detection system (PerkinElmer) as per manufacturer’s instructions. β-ACTIN was used as loading control.

### Immunoprecipitation Assay

Immunoprecipitation assays were carried out following a modified version of the protocol provided by Millipore, USA. In brief, macrophages were gently resuspended and lysed in ice-cold RIPA buffer. The cell lysates obtained were subjected to pre-clearing with BSA-blocked Protein A beads. The amount of protein was estimated in the supernatant and equal amount of protein was incubated with IgG or anti-SREBP2 antibody for 4 h at 4 °C. The immune complexes were captured on protein A agarose beads (Bangalore Genei, India) at 4 °C for 4 h. The beads were separated, washed and boiled in Laemmli buffer for 10 min. These bead free samples were analyzed for respective target molecules by immunoblotting. Light chain specific secondary antibody was used for immunoblotting after immunoprecipitation.

### Chromatin Immunoprecipitation (ChIP) Assay

ChIP assays were carried out using a protocol provided by Upstate Biotechnology and Sigma-Aldrich with certain modifications. Briefly, macrophages were fixed with 3.6 % formaldehyde for 15 min at room temperature followed by inactivation of formaldehyde with addition of 125 mM glycine for 10 min. Nuclei were lysed in 0.1 % SDS lysis buffer [50 mM Tris-HCl (pH 8.0), 200 mM NaCl, 10 mM HEPES (pH 6.5), 0.1 % SDS, 10 mM EDTA, 0.5 mM EGTA, 1 mM PMSF, 1 μg/ml of each aprotinin, leupeptin, pepstatin, 1 mM Na_3_VO_4_ and 1 mM NaF]. Chromatin was sheared using Bioruptor Plus (Diagenode, Belgium) at high power for 70 rounds of 30 sec pulse ON and 45 sec pulse OFF. Chromatin extracts containing DNA fragments with an average size of 500 bp were immunoprecipitated with SIRT6 or G9a or H3K9Ac or H3K9me1 or H3K9me2 or β-CATENIN antibodies or rabbit preimmune sera complexed with Protein A agarose beads (Bangalore Genei, India). Immunoprecipitated complexes were sequentially washed with Wash Buffer A, B and TE [Wash Buffer A: 50 mM Tris-HCl (pH 8.0), 500 mM NaCl, 1 mM EDTA, 1 % Triton X-100, 0.1 % Sodium deoxycholate, 0.1 % SDS and protease/phosphatase inhibitors; Wash Buffer B: 50 mM Tris-HCl (pH 8.0), 1 mM EDTA, 250 mM LiCl, 0.5% NP-40, 0.5 % Sodium deoxycholate and protease/phosphatase inhibitors; TE: 10 mM Tris-HCl (pH 8.0), 1 mM EDTA] and eluted in elution buffer [1 % SDS, 0.1 M NaHCO3]. After treating the eluted samples with RNase A and Proteinase K, DNA was precipitated using phenol-chloroform-ethanol method. Purified DNA was analyzed by quantitative real time RT-PCR. All values in the test samples were normalized to amplification of the specific gene in Input and IgG pull down and represented as fold change in modification or enrichment. All ChIP experiments were repeated at least three times. The list of primers is detailed in Supplementary Table 3.

### Sequential ChIP Assay

The protocol for sequential ChIP was adopted from (de Medeiros, 2011; Truax and Greer, 2012). Briefly, the DNA fragments obtained following sonication [in lysis buffer; 1 % SDS, 10 mM EDTA, 50 mM Tris-HCl (pH 8.0)] were immunoprecipitated with SREBP2-complexed Protein A beads. After first pull down, beads were washed with Re-ChIP Buffer [2 mM EDTA, 500 mM NaCl, 0.1 % SDS, 1 % NP40], followed by elution of DNA in Re-ChIP elution buffer [2 % SDS, 15 mM DTT in TE] at 37 °C for 30 min. The eluted DNA was subjected to subsequent round to immunoprecipitation with Protein-A beads pre-complexed with G9a or rabbit pre-immune sera. Immunoprecipitated complexes were sequentially washed with Wash Buffer A, B and TE [Wash Buffer A: 20 mM Tris-HCl (pH 8.0), 150 mM NaCl, 2 mM EDTA, 1% Triton X-100, 0.1% SDS and protease/phosphatase inhibitors; Wash Buffer B: 20 mM Tris-HCl (pH 8.0), 2 mM EDTA, 500 mM NaCl, 1 % Triton X-100, 0.1 % SDS and protease/phosphatase inhibitors; Wash Buffer C: 10 mM Tris-HCl (pH 8.0), 1 mM EDTA, 1 % sodium deoxycholate, 1 % NP40, 250 mM LiCl and protease/phosphatase inhibitors; TE: 10 mM Tris-HCl (pH 8.0), 1 mM EDTA and protease/phosphatase inhibitors] and eluted [0.1 M NaHCO_3_, 1 % SDS], purified and subjected to qRT-PCR (as described previously). The fold change of SREBP2-G9a versus SREBP2-IgG upon infection signified the co-occupancy of the two factors at concerned promoters. The list of primers is given in Supplementary Table 3.

### Isolation and culture of murine bone marrow derived macrophages

Mice tibia and femur were flushed with ice-cold DMEM containing 10 % fetal bovine serum from WT (littermate control) and *sirt6* KO mice. Bone marrow was collected in 50 ml tube and bone marrow clusters were disintegrated by vigorous pipetting. The cell suspension was centrifuged at 1500 rpm for 5 min at 4 °C followed by two washes with DMEM containing 10 % fetal bovine serum. Then the cells were suspended in DMEM containing 10 % fetal bovine serum and 20 % of L929 cell supernatant and seeded at 1 million cells per well and incubated at 37 °C, 5 % CO_2_ and 95 % humidity in a CO_2_ incubator. The medium was supplemented on the 3^rd^ and 5^th^ day with DMEM containing 10 % fetal bovine serum and 20 % L929 cell supernatant. Post 7 days of differentiation, the cells were used for further experiments.

### *In vitro* Mtb CFU

BALB/c peritoneal macrophages transfected with *Ehmt2*, *Sirt6*, Srebf2, cholesterol genes (*Lrp2, Mvd, Aacs, Hmgcs, Dhcr24*), *Nfe2l2*, *Ctnnb1* or non-targeting siRNA for 24 h; or BMDMs obtained from *Sirt6* KO mice were infected with Mtb H37Rv at MOI 5 for 4 h. Post 4 h, the cells were thoroughly washed with PBS to remove any surface adhered bacteria and medium containing amikacin (0.2 mg/ml) was added for 2 h to kill any extracellular mycobacteria. After amikacin treatment, the cells thoroughly washed with PBS were taken as 0 h time point and a duplicate set was maintained in antibiotic free medium for next 48 h. Intracellular mycobacteria was enumerated by lysing macrophages with 0.06 % SDS in 7H9 Middlebrook medium. Appropriate dilutions were plated onto Middlebrook 7H11 agar plates supplemented with OADC (oleic acid, albumin, dextrose, catalase). Total colony forming units (CFUs) were counted after 21 days of plating.

### Microtomy and Hematoxylin and Eosin (H&E) staining

Microtome sections (5 μm) were obtained from formalin-fixed, paraffin-embedded mouse lung tissue samples using Leica RM2245 microtome. These sections were first deparaffinized and rehydrated. The rehydrated sections were subjected to Hematoxylin staining followed by Eosin staining as per manufacturer instructions. The sections were then dehydrated and mounted with coverslip using permount. Sections were given to consultant pathologist for blinded analyses.

### Cryosection preparation

The excised and fixed lungs were placed in the optimal cutting temperature (OCT) media (Jung, Leica). Cryosections of 10-15 µm were prepared using Leica CM 1510 S or Leica CM 3050 S cryostat with the tissue embedded in OCT being sectioned onto glass slides and then stored at −80 °C.

### Immunofluorescence (IF)

Treated/infected macrophages were fixed with 3.6 % formaldehyde for 30 min at room temperature. The cells were washed with PBS and blocked in 2 % BSA in PBST. After blocking, cells were stained with LRP2 overnight at 4 °C. Then they were incubated with DyLight 488-conjugated secondary antibody for 2 h and nuclei were stained with DAPI. The coverslips were mounted on a slide with glycerol. For IF of the cryosections, frozen sections were thawed to room temperature. After blocking with 2 % BSA containing saponin, the sections were stained with specific antibodies overnight at 4 °C. The sections were then incubated with DyLight 488-conjugated secondary antibody for 2 h and nuclei were stained with DAPI. A coverslip was mounted on the section with glycerol as the medium. Confocal images were taken with Zeiss LSM 710 Meta confocal laser scanning microscope (Carl Zeiss AG, Germany) using a plan-Apochromat 63X/1.4 Oil DIC objective (Carl Zeiss AG, Germany) and images were analyzed using ZEN black software. CTCF (corrected total cell fluorescence) was calculated as (fluorescence observed in an area of a cell – fluorescence of background for the same area) using ImageJ. Cells boundaries were demarcated based on brightfield image and the fluorescence intensities of different channels were measured. Background fluorescence intensity was measured from a field devoid of cells.

### Filipin fluorescence staining for Free Cholesterol

Filipin complex (Sigma-Aldrich, USA) was utilized to assess free cholesterol following protocol from (Leventhal et al, 2001). Briefly, mouse peritoneal macrophages were fixed with 3.6 % paraformaldehyde for 1 h at room temperature. After incubation, cells were washed with 1X PBS followed by incubation in 1.5 mg glycine per ml PBS for 10 min at room temperature. Filipin staining was then performed at a final concentration of 0.05 mg/ml in PBS for 2 h at room temperature. Cell were washed thrice with 1X PBS and nuclei were stained with propidium iodide (PI). For Filipin staining of cryosections, frozen sections were thawed to room temperature. After blocking with 2 % BSA containing saponin, the sections were stained with Filipin (0.05 mg/ml in PBS) and PE-conjugated F4/80 (macrophage marker) for 2 h at room temperature. The samples were mounted on glycerol. Images were captured in Zeiss LSM 710 confocal laser scanning microscope as described above.

### CellROX Oxidative Stress Reagent staining

CellROX Deep Red Reagent (Thermo Fisher Scientific, USA) was utilized to measure oxidative stress in macrophages as per manufacturer’s instructions. In brief, siRNA transfected mouse peritoneal macrophages were treated with CellROX Deep Red Reagent at a final concentration of 5 μM and incubated for 30 min at 37 °C. Cells were then washed with 1X PBS thrice followed by fixation with 3.6 % formaldehyde for 15 min. Nuclei were stained with DAPI and images were captured in Zeiss LSM 710 confocal laser scanning microscope.

## Supplementary Tables

**Supplementary Table 1:**
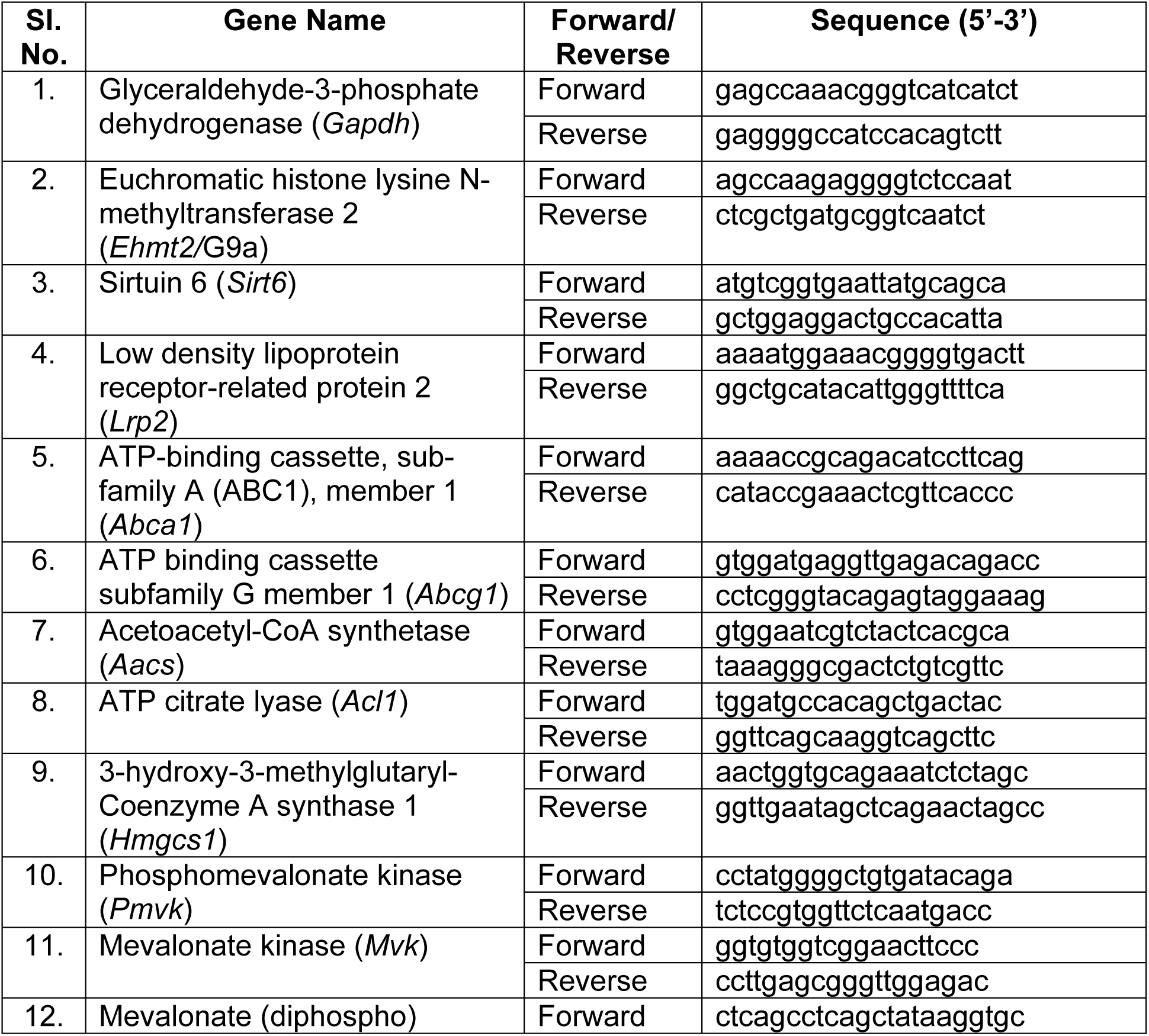
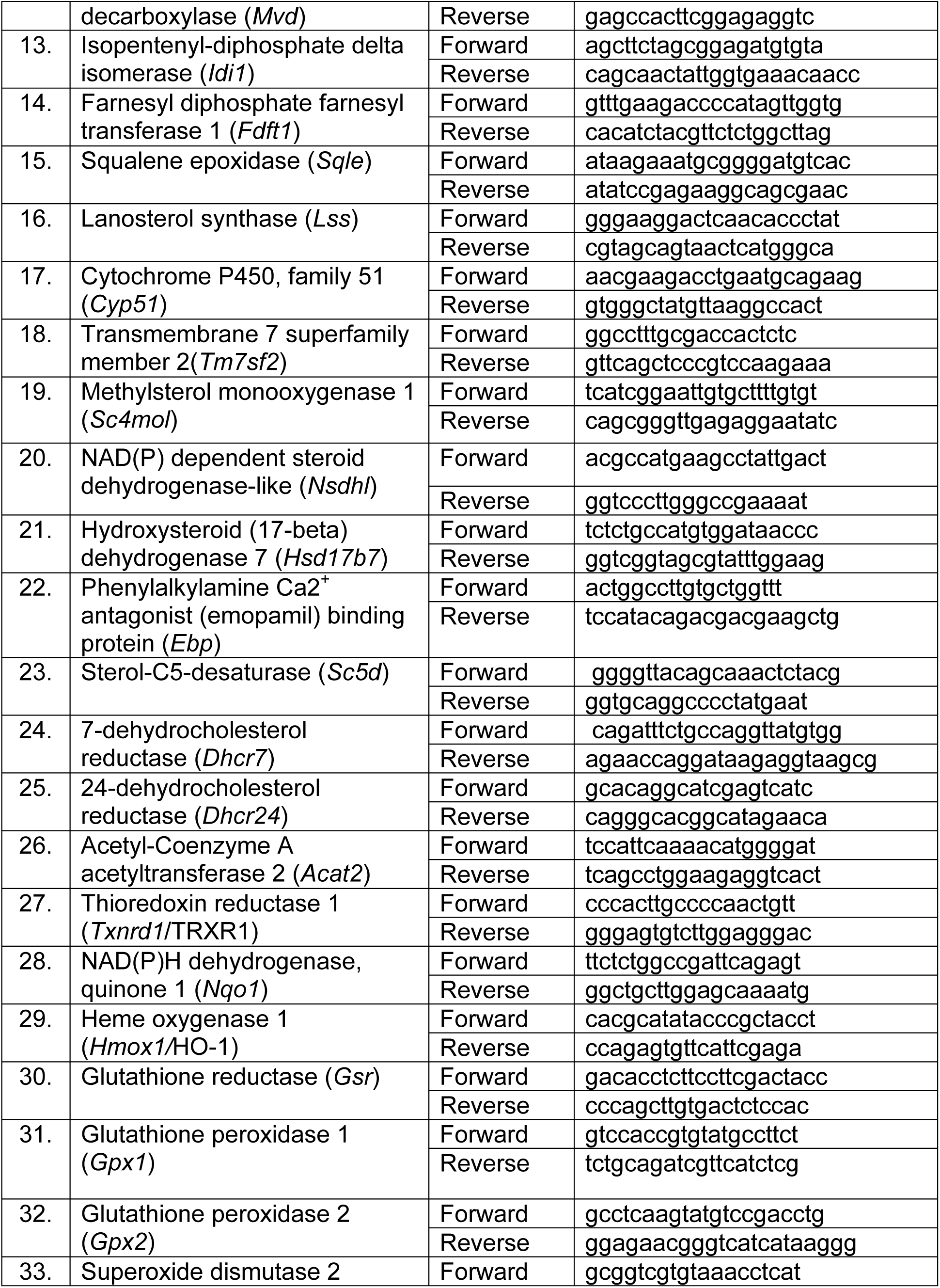
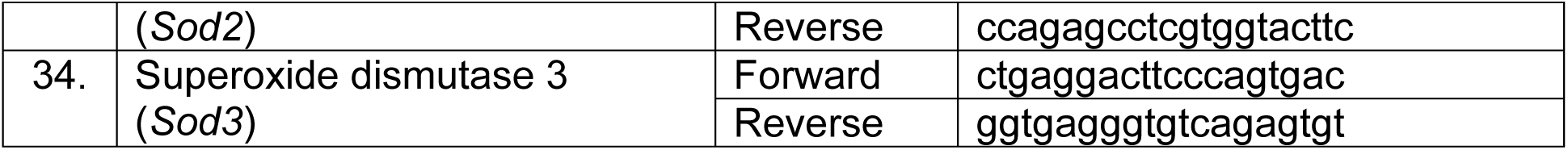
List of primers for mouse gene expression analyses.

**Supplementary Table 2:**
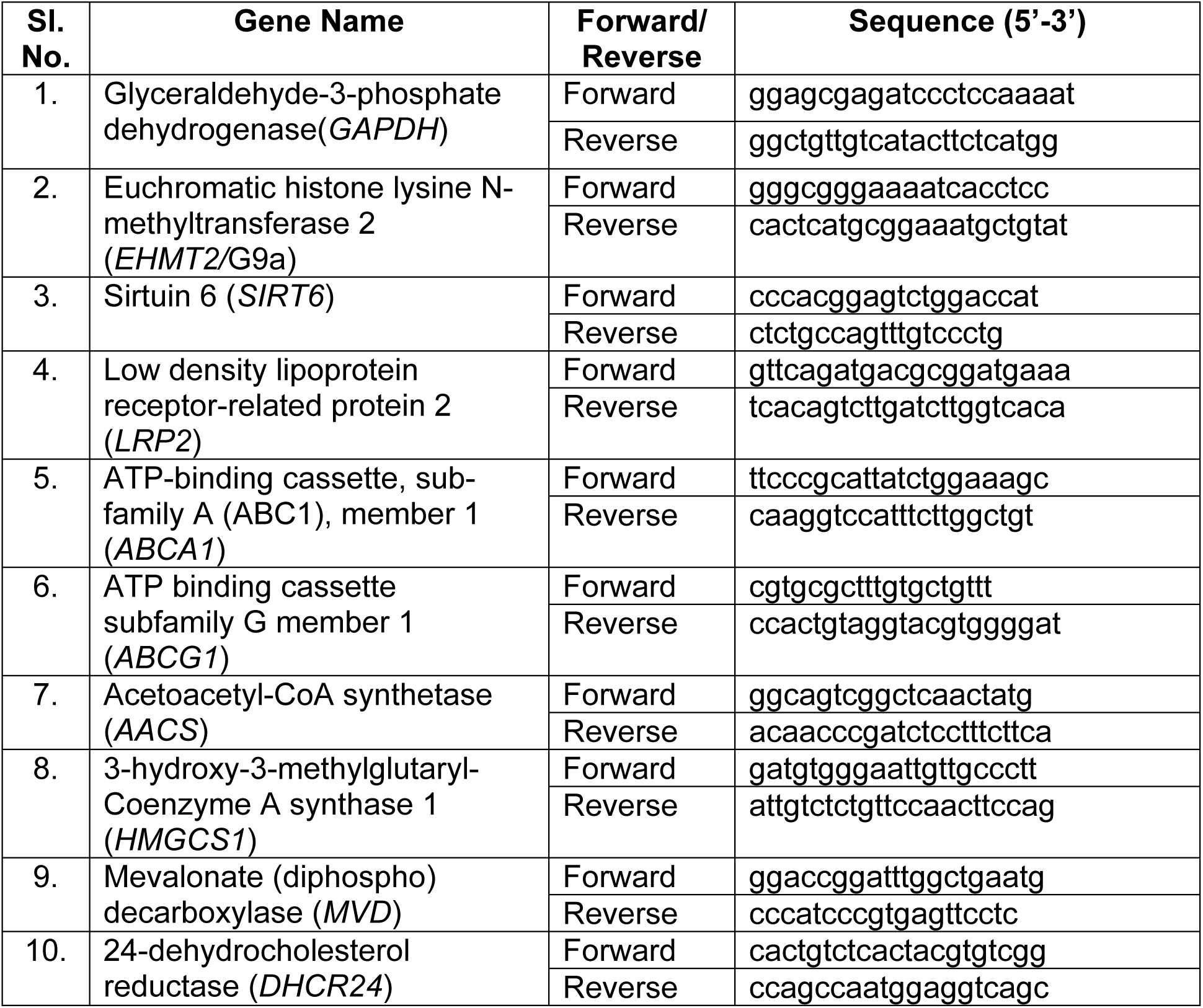
List of primers for human gene expression analyses.

**Supplementary Table 3:**
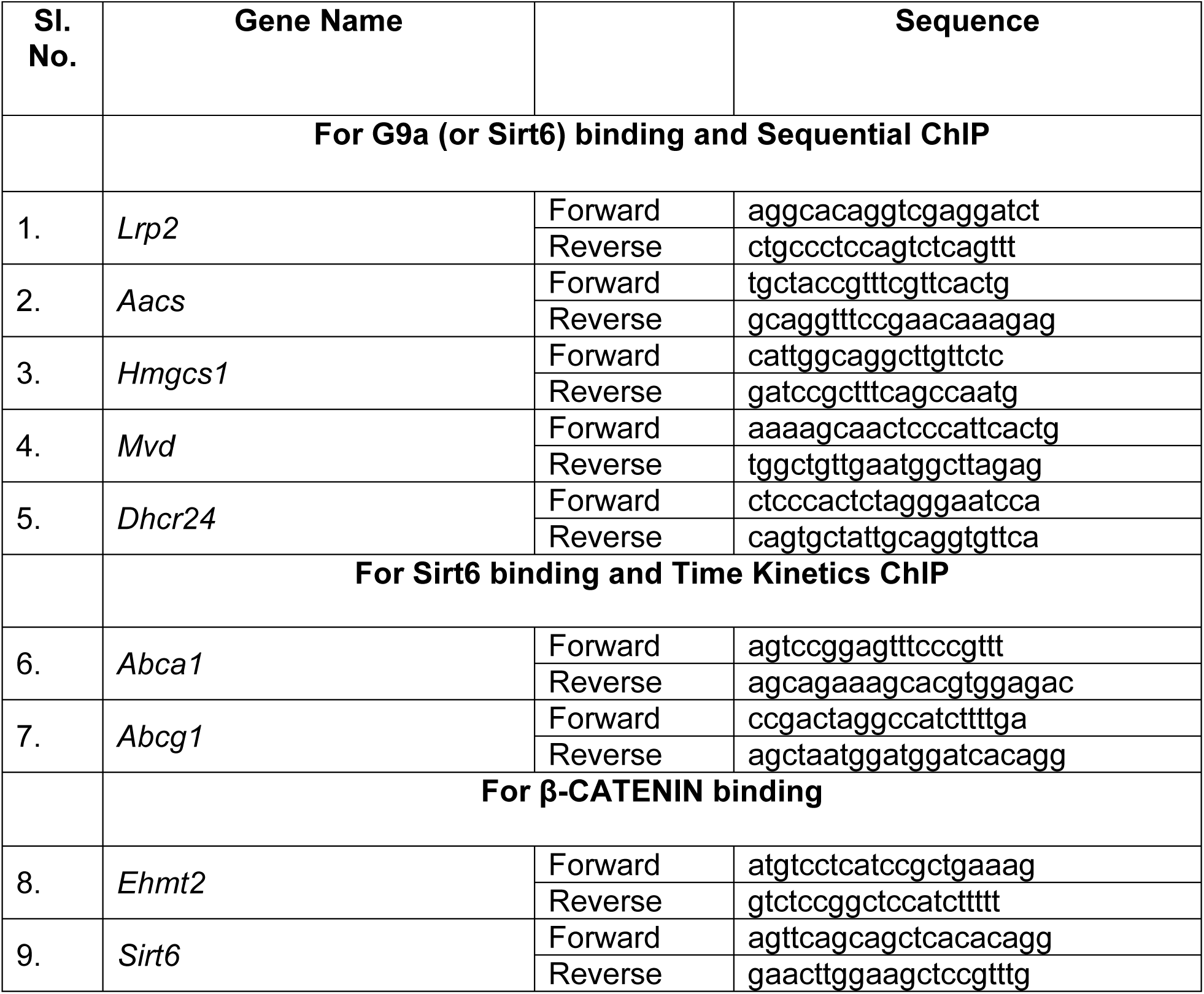
List of primers for ChIP assays.

